# The phosphodiesterase 2A regulates lymphatic endothelial development via cGMP-mediated control of Notch signaling

**DOI:** 10.1101/2023.01.18.524585

**Authors:** Claudia Carlantoni, Leon Liekfeld, Sandra A. Hemkemeyer, Danny Schreier, Ceren Saygi, Roberta Kurelic, Silvia Cardarelli, Joanna Kalucka, Christian Schulte, Manu Beerens, Reiner Mailer, Tilman Schäffer, Fabio Naro, Manuela Pellegrini, Viacheslav O. Nikolaev, Thomas Renné, Maike Frye

## Abstract

During vascular development endothelial junctions mature and vessel integrity is established to form the endothelial barrier. The molecular mechanisms by which lymphatic vessels induce cell contact inhibition are not understood. Here, we uncover the cGMP-dependent phosphodiesterase 2A (PDE2A) as a selective regulator of lymphatic, but not blood endothelial contact inhibition. Conditional deletion of *Pde2a* in mouse embryos reveals severe lymphatic dysplasia, while large blood vessel architecture remains unaltered. In the absence of PDE2A, human lymphatic endothelial cells fail to induce mature junctions and cell cycle arrest, while cGMP levels, but not cAMP levels, are increased. Loss of PDE2A-mediated cGMP hydrolysis leads to downregulation of NOTCH signaling. Vice versa, DLL4-induced NOTCH activation restores junctional maturation in PDE2A-deficient lymphatic endothelial cells. Our data demonstrate that PDE2A selectively modulates a crosstalk between cGMP and NOTCH signaling to finetune lymphatic development and suggest that PDE2A may be a druggable target to control lymphatic leakage and regeneration.

## Introduction

At the onset of lymphatic and blood vascular development dynamic remodeling of endothelial cell-cell contacts is the prerequisite for initial sprout formation (Zhang et al., 2020a, Szymborska and Gerhardt, 2018). At later stages cell junctions mature and vessel integrity is established to form the endothelial barrier between the circulating lymph or blood and the surrounding tissue (Potente and Mäkinen, 2017).

Endothelial cells (ECs) exit their cell cycle and undergo cell contact inhibition when they reach confluency. In blood ECs (BECs) it has been suggested that the formation of stable junctions induces intracellular signals to restrains the capacity of the cells to respond to proliferative signals (Choi et al., 2015). Furthermore, Vascular endothelial (VE)-cadherin-mediated signaling (Lampugnani et al., 2003, Baumeister et al., 2005), YES-associated protein 1 (YAP) (Ritchey et al., 2019) and NOTCH signaling (Noseda et al., 2004) have been shown to control BEC contact inhibition. While this process has been extensively studied in BECs, little is known about the molecular mechanisms by which contact inhibition is induced during development of the lymphatic vasculature, despite its key role in fluid homeostasis, fat absorption, immune cell transport and in diseases, such as lymphedema, cancer and immunological dysfunction (Oliver et al., 2020).

Dynamics of the endothelial barrier are further controlled by the combined action of intracellular mechanisms, such as TIE, NOTCH, FOCX2, GTPase and cyclic nucleotide signaling (Stritt et al., 2021, Claesson-Welsh et al., 2021, Sabine et al., 2015, Surapisitchat and Beavo, 2011), and extracellular factors, such as fluid mechanics and matrix stiffness (Dorland and Huveneers, 2017, Gordon et al., 2020). To this end, lymphatic ECs (LECs) and BECs are equipped with a plethora of similar molecules, which can have multifaceted roles in different EC types and developmental stages. For example, deletion of VE-cadherin (*Cdh5*) is dispensable for junctional maintenance of mature blood vessels in mouse skin and brain (Frye et al., 2015), mature mouse dermal lymphatic vessels (Hägerling et al., 2018) and cell-cell junctions of human dermal LECs (Frye et al., 2020). Recent findings further showed that the receptor tyrosine kinase EphB4 selectively regulates junctional integrity of postnatal and adult lymphatics (Frye et al., 2020) and adult cardiac capillaries (Luxán et al., 2019).

Cyclic nucleotide phosphodiesterases (PDEs) comprise a superfamily of metallophosphohydrolases that specifically hydrolyze the second messenger cyclic adenosine monophosphate (cAMP) and/or cyclic guanosine monophosphate (cGMP) to AMP and GMP, respectively. In mammals, PDEs can be divided in three groups depending on their substrate selectivity: cAMP-specific, cGMP-specific, and PDEs with dual specificity. Among the latter category, PDE2 exhibits dual substrate specificity for cAMP and cGMP and is the only cAMP-hydrolyzing PDE, which is allosterically activated by cGMP. Due to this unique role, PDE2 has been suggested to play a role at the center of the crosstalk between cGMP and cAMP signaling (Weber et al., 2017). In BECs, relative expression levels of PDE2A and another PDE, the cGMP-inhibited PDE3A, have been shown to regulate blood endothelial junctions (Surapisitchat et al., 2007, Chen et al., 2016), but the function of PDE2A in LECs is not known. Here, we studied the role of the phosphodiesterase PDE2A in lymphatic development and junctional integrity using conditional *Cre/loxP* mediated gene deletion in mice and corresponding analysis in primary human LECs. We found that mouse embryos developed severe lymphatic dysplasia upon deletion of endothelial *Pde2a*, while large blood vessels were unaltered. In the absence of PDE2A, human LECs failed to induce mature CLDN5^High^ junctions and cell cycle arrest. Consistent with defective junctional maturation, loss of endothelial *Pde2a* expression in mice interfered with lymphatic maturation and blocked contact inhibition at later stages during embryonic development. Unexpectedly, lymphatic cGMP levels, but not cAMP levels, were increased in PDE2A-deficient LECs. Intriguingly, dysregulated lymphatic cGMP was associated with reduced NOTCH signaling, while exogenous activation of NOTCH via its ligand Delta-like 4 (DLL4) was able to rescue CLDN5^High^ expression and junctional maturation. Our data uncover a selective mechanism of lymphatic maturation via the previously unappreciated PDE2A/cGMP/NOTCH axis.

## Results

### Expression of the phosphodiesterase 2A is enriched in lymphatic endothelial cells

During embryonic development, lymphatic endothelial progenitors and venous ECs within the cardinal vein experience extracellular matrix (ECM) stiffness around 4 kPa (Frye et al., 2018). They are tightly attached to the underlying basement membrane and show a flattened, spread-out monolayer morphology, while in a soft microenvironment (0.2 kPa) lymphatic endothelial progenitors preferably migrate (Frye et al., 2018). Similarly, human LECs in culture switch from a migratory, spindle shape phenotype on very soft substrates (0.1 kPa) to a flattened morphology with the capacity to form a continuous monolayer on moderate stiffness substrates (4 kPa) (**Figure 1A**).

**Figure 1:**
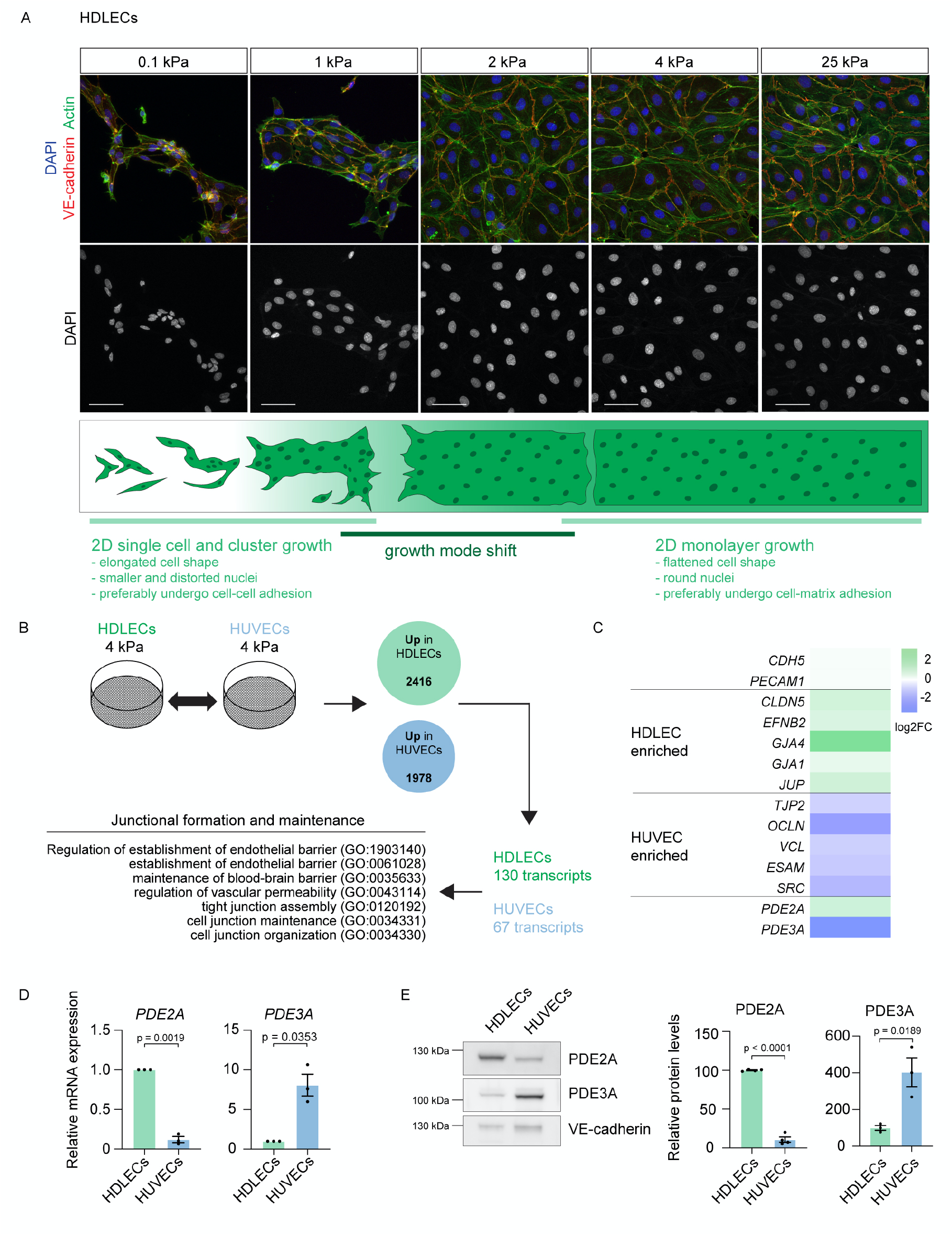
PDE2A is enriched in lymphatic endothelial cells. **A)** Representative immunofluorescence staining of human dermal lymphatic endothelial cells (HDLECs), grown on polyacrylamide stiffness substrates ranging from 0.1 to 25 kPa, using an antibody against VE-cadherin (red), phalloidin (green, to visualize Actin) and DAPI (blue). Single channel images for DAPI are shown. Scale bar: 50 μm. **B)** Global transcriptome analysis of HDLECs and HUVECs grown on physiological 4 kPa stiffness substrates. Number of enriched transcripts (n = 3 biological replicates) in HDLECs and human umbilical vein endothelial cells (HUVECs) are shown. Among these, differentially regulated transcripts associated with gene ontology (GO) terms of junctional formation and maintenance were selected. **C)** Heatmap of selected enriched transcripts in HDLECs and HUVECs. **D)** Relative mRNA expression levels of *PDE2A* and *PDE3A* in HDLECs and HUVECs from n = 3 independent experiments. Mean ± s.e.m., p-value: One-sample *t*-test. **E)** Western Blot analysis and quantification (n = 3 independent experiments) of PDE2A and PDE3A expression in HDLECs and HUVECs. VE-cadherin was used as endothelial loading control. Mean ± s.e.m., p-value: Unpaired Student’s *t*-test.

To identify potential selective regulators of lymphatic and blood EC junction formation and vessel maturation in a physiological stiffness environment, we performed global transcriptome analysis (RNAseq) of lymphatic (human dermal lymphatic ECs, HDLECs) and venous ECs (human umbilical vein ECs, HUVECs) grown on junction-reinforcing 4 kPa stiffness substrates. RNAseq analysis revealed cell type specific upregulation of 2416 and 1978 transcripts in HDLECs and HUVECs, respectively (log2 fold change > 0.5 or < −0.5) (**Figure 1B**). Within these upregulated transcripts we identified 130 HDLEC transcripts and 67 HUVEC transcripts that were annotated to gene ontology (GO) terms related to endothelial junction formation and maintenance (**Figure 1B**).

As expected, the junctional molecules VE-cadherin *(CDH5*) and CD31 (*PECAM1*) were not differentially expressed between both cell types. However, mRNA levels of CLDN5 (*CLDN5*) and EphrinB2 (*EFNB2*), which we have previously identified to regulate LEC junctions (Frye et al., 2020), were upregulated in HDLECs (**Figure 1C**). Notably, strongly enriched transcripts included the PDEs *PDE2A* (upregulated in HDLECs) versus *PDE3A* (upregulated in HUVECs) (**Figure 1C**) of which the relative expression levels have been shown to regulate blood endothelial permeability (Surapisitchat et al., 2007). Quantitative real time (qRT)-PCR analysis of HDLECs and HUVECs grown for 72 h confirmed RNAseq findings and showed an 8.3-fold decrease of *PDE2A* and an 8.1-fold increase of *PDE3A* mRNA levels in HUVECs compared to HDLECs (**Figure 1D**).

Similarly, PDE2A and PDE3A showed inverse protein expression levels in HDLECs compared to HUVECs (**Figure 1E**). We therefore sought to analyze if enriched expression of PDE2A in LECs could reflect a unique role of this PDE in the regulation of lymphatic junctions compared to venous blood endothelial junctions.

### Endothelial-specific deletion of Pde2a results in severe dysplasia of the embryonic lymphatic vasculature

Global deletion of *Pde2a* (*Pde2a*^*-/-*^) however results in embryonic lethality around embryonic day (E) 14.5-15.5, which was suggested to be caused by anemia, hemorrhages and reduced fetal liver size (Assenza et al., 2018, Barbagallo et al., 2020, Stephenson et al., 2012). Consistent with our human data, single cell (sc)RNAseq data from the Tabula muris database (Schaum et al., 2018) show enrichment of *Pde2a* gene expression in LECs. Using literature-curated marker genes of cell phenotypes, we exemplarily identified an LEC cluster in mouse lungs (**Supplemental figure 1A)**, which showed high expression of *Pde2a* in LECs and low expression of *Pde3a* in pulmonary ECs with a main presence in non-EC types (**Supplemental figure 1B, C**). We therefore focused on the analysis of PDE2A function in LECs *in vivo*.

In order to investigate the role of PDE2A in LECs and BECs, we ablated *Pde2a* expression in BECs prior to LEC commitment using the *Tie2-Cre* mice in combination with a newly generated floxed *Pde2a* allele (**Figure 2A, Supplemental figure 2A**,**B**). Efficient deletion of endothelial *Pde2a* was demonstrated by RT-PCR of E14.5 sorted CD31^+^;PDPN^+^ LECs (77%) and CD31^+^; PDPN^-^ BECs (91%) (**Figure 2B**). Analysis of immunostained transverse vibratome sections of E13.5 and E14.5 embryos revealed Prospero Homeobox 1 (PROX1)^+^ primordial thoracic ducts (pTD) and peripheral longitudinal lymphatic vessels (PLLV) (commonly referred to as ‘jugular lymph sac’) that were massively enlarged in *Pde2a*^*flox/flox*^; *Tie2-Cre* embryos in comparison to Cre-negative control embryos (**Figure 2C, Supplemental figure 3A**). Jugular lymph sacs of heterozygous *Pde2a*^*flox/+*^; *Tie2-Cre* were indistinguishable from Cre-negative control embryos (**Supplemental figure 3B**).

**Figure 2:**
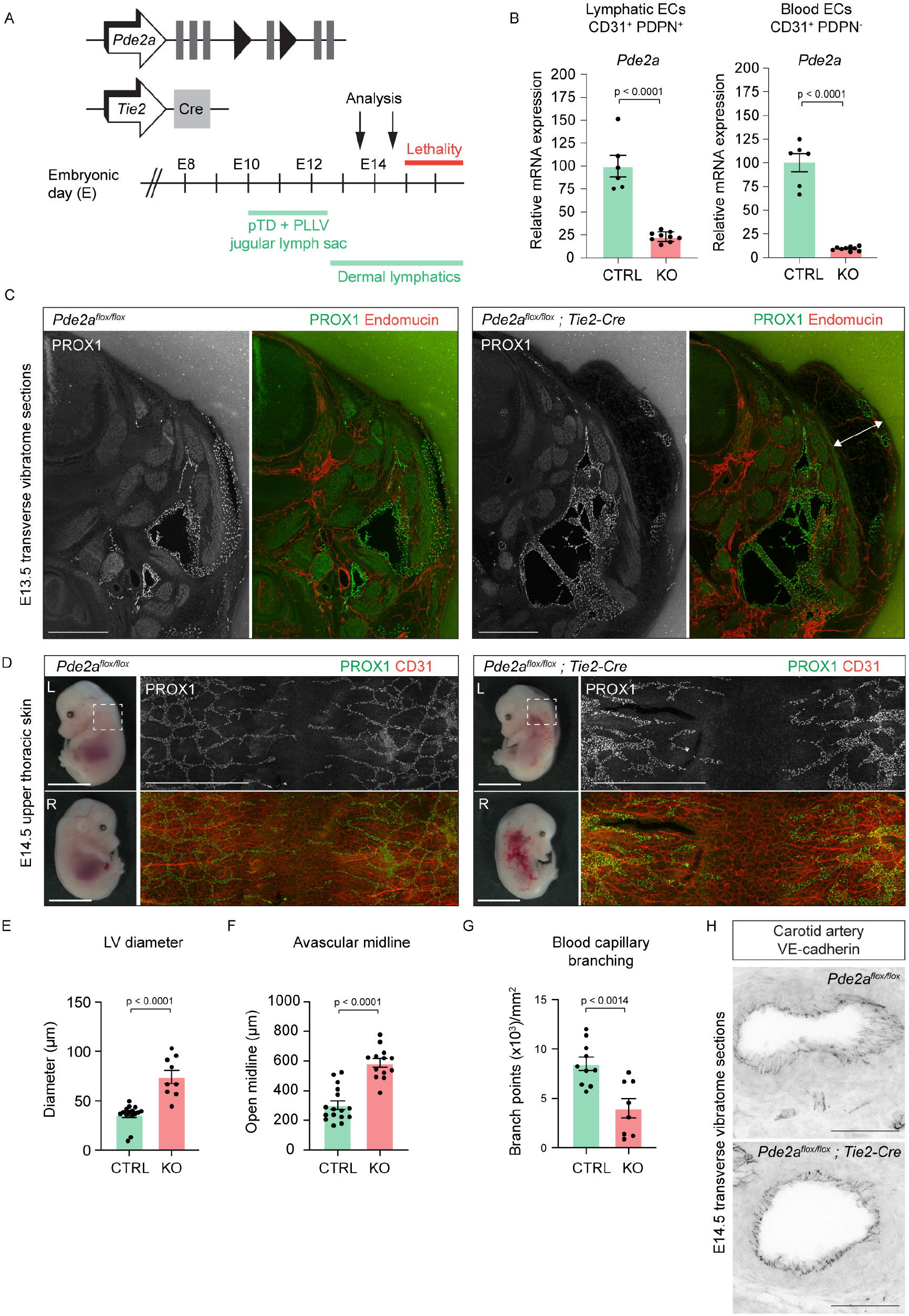
Endothelial *Pde2a* deletion in the *Tie2*-lineage impairs embryonic lymphatic development. **A)** Schematic of the genetic constructs and analyzed embryonic stages (E). **B)** Relative mRNA expression of *Pde2a* in LECs and BECs sorted from E14.5 back skins of n = 6 *Pde2a*^*flox/flox*^; *Tie2-Cre* embryos (KO) and n = 9 *Pde2a*^*flox/flox*^ littermate embryos (CTRL). Mean ± s.e.m., p-value: Unpaired Student’s *t*-test. **C)** Representative immunofluorescence staining of 100μm transverse vibratome sections from E13.5 KO and CTRL embryos stained with antibodies against PROX1 (green) and Endomucin (red). Single channel images for PROX1 are shown. KO embryos display enlarged jugular lymphatic structures and nuchal edema (arrow). Scale bar: 500 μm. **D)** Representative brightfield images of E14.5 *Pde2a* KO and CTRL embryos and immunofluorescence staining of whole-mount upper thoracic back skins stained with antibodies against PROX1 (green) and CD31 (red). Single channel images for PROX1 are shown. Scale bar: 500 μm. **E**,**F)** Quantification of lymphatic vessel (LV) diameter (**E**) and avascular midline (**F**) in E14.5 upper thoracic back skins of n = 9-13 KO and n = 15-16 CTRL embryos. Mean ± s.e.m., p-value: Unpaired Student’s *t*-test. **G)** Quantification of blood capillary branching in E14.5 upper thoracic back skins of n = 8 KO and n = 10 CTRL embryos. Mean ± s.e.m., p-value: Unpaired Student’s *t*-test. **H)** Representative immunofluorescence staining of the carotid artery in 100μm transverse vibratome sections of E14.5 KO and CTRL embryos stained with an antibody against VE-cadherin. Notably, carotid artery morphology and VE-cadherin^+^ junctions were not altered in KO embryos. Scale bar: 50 μm.

**Figure 3:**
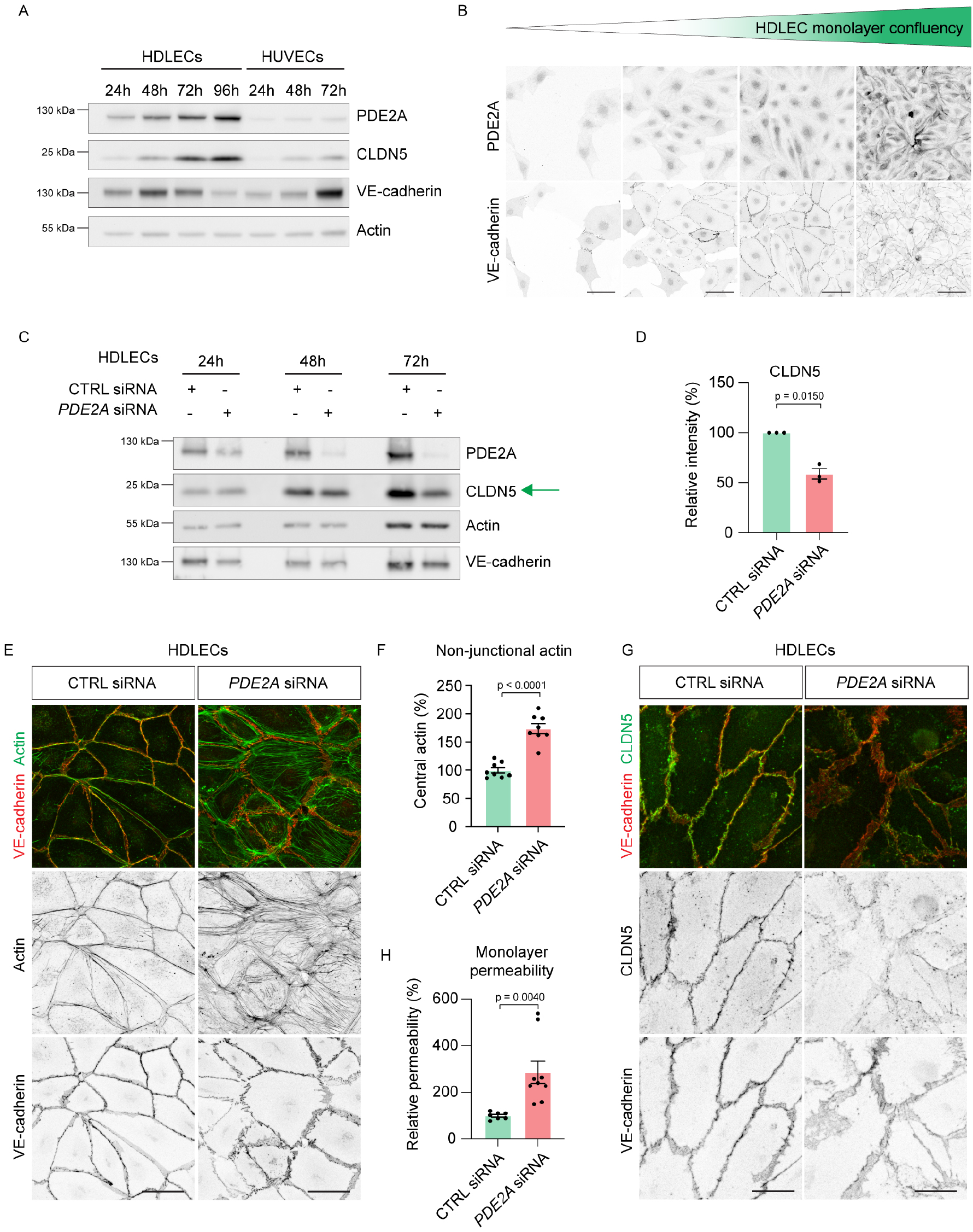
Lymphatic junctions fail to mature in the absence of PDE2A. **A)** Western Blot analysis of HDLECs and HUVECs cultured from 24 to 96h and 72h, respectively. Lysates were probed with antibodies against PDE2A, CLDN5, VE-cadherin and Actin as loading control. **B)** Representative immunofluorescence staining of HDLECs cultured from 24 to 96h and stained with antibodies against PDE2A and VE-cadherin. With increasing monolayer confluency PDE2A expression is increased. Scale bar: 50 μm. **C)** Western Blot analysis of CTRL and *PDE2A* siRNA treated HDLECs cultured for 24, 48 and 72h. Lysates were probed with antibodies against PDE2A, CLDN5, VE-cadherin and Actin as loading control. Notably, in the absence of PDE2A, HDLECs fail to form CLDN5^High^ junctions (green arrow). **D)** Quantification of CLDN5 protein expression in 72h CTRL and *PDE2A* siRNA treated HDLECs from n = 3 independent experiments. Mean ± s.e.m., p-value: One-sample *t*-test. **E)** Representative immunofluorescence staining of 72h CTRL and *PDE2A* siRNA treated HDLECs stained with an antibody against VE-cadherin (red) and phalloidin (green, to visualize Actin). Single channel images of VE-cadherin and phalloidin are shown. Scale bar: 50 μm. **F)** Quantification of relative central actin (total actin – junctional actin) from 72h CTRL and *PDE2A* siRNA treated HDLECs from n = 2 independent experiments. Mean ± s.e.m., p-value: Unpaired Student’s *t*-test. **G)** Representative immunofluorescence staining of 72h CTRL and *PDE2A* siRNA treated HDLECs using antibodies against VE-cadherin (red) and CLDN5 (green). Single channel images are shown. Scale bar: 20 μm. **H)** Quantification of monolayer permeability to 40kDa FITC-dextran in CTRL and *PDE2A* siRNA treated HDLECs from n = 2 independent experiments, mean ± s.e.m., p-value: Unpaired Student’s *t*-test.

Furthermore, *Pde2a*^*flox/flox*^; *Tie2-Cre* embryos exhibited large nuchal edema (**Figure 2C**) that can be indicative of a dysfunctional lymphatic vasculature (Burger et al., 2015). Consistent with these findings, wholemount immunostaining of E14.5 upper thoracic back skins revealed disturbed lymphangiogenesis with disconnected and enlarged lymphatic vessels in *Pde2a*^*flox/flox*^; *Tie2-Cre* embryos (**Figure 2D-F**). A similar lymphatic phenotype was observed in NRP2^+^ dermal lymphatic vessels of global E14.5 *Pde2a*^*-/-*^ mutant embryos compared to littermate control embryos (**Supplemental figure 3C**).

Blood capillary branching was reduced in the midline region of *Pde2a*^*flox/flox*^; *Tie2-Cre* embryos (**Figure 2G**), while VE-cadherin^+^ E14.5 carotid artery diameter (**Figure 2H**) and overall large blood vessel morphology was normal (Endomucin^+^ vessels in **Figure 2C** and CD31^+^ larger vessels in **Figure 2D**).

Taken together, these results indicate that PDE2A plays an important role in lymphatic development.

### In the absence of PDE2A expression lymphatic junctions fail to mature in vitro

We next sought to analyze if loss of PDE2A could contribute to lymphatic vessel dysplasia through dysregulation of lymphatic junctions. HDLECs were cultured for 24h, 48h, 72h and 96h to correlate changes in protein levels with increasing lymphatic monolayer confluency. As expected, protein expression of CLDN5, a major lymphatic junctional molecule (Frye et al., 2020), was induced with increasing monolayer confluency, demonstrating the formation of stable and mature lymphatic junctions (**Figure 3A**). Notably, VE-cadherin was highly expressed in immature lymphatic junctions but was downregulated with increasing CLDN5 protein expression.

Interestingly, PDE2A protein expression also increased with lymphatic monolayer maturation (**Figure 3A**). Immunofluorescence staining using antibodies against PDE2A and VE-cadherin further confirmed immunoprint data (**Figure 3B**). In contrast, HUVECs cultured for 24h, 48h and 72h showed an expected strong increase in VE-cadherin and a moderate increase in CLDN5 protein signal, however low PDE2A levels was not altered (**Figure 3A**).

To assess the functional consequence of PDE2A loss, we performed siRNA-mediated knockdown of *PDE2A* expression in HDLECs with increasing monolayer confluency states. Interestingly, loss of PDE2A only resulted in decreased CLDN5 expression levels **(Figure 3C**, green arrow, **3D**) in stable lymphatic monolayers (72h culture) having high PDE2A protein levels (**Figure 3A**), Together the data indicate that *PDE2A*-deficient LECs fail to form mature and stable CLDN5^+^ junctions *in vitro*.

Upon siRNA-mediated knockdown of *PDE2A* expression, immunofluorescence analysis of VE-cadherin, actin, and CLDN5 further showed disruption of 72h cultured HDLECs but not HUVECs (**Supplemental figure 4**), characterized by decreased cortical actin (**Figure 3E,F**) and dispersed junctional CLDN5 (**Figure 3G**), Consistent with these morphological findings, deletion of *PDE2A* impaired barrier function and led to increased HDLEC monolayer permeability (**Figure 3H)**. Conclusively, in contrast to venous blood ECs, PDE2A expression is strongly upregulated in LECs during the formation of stable cell-cell junctions. Conversely, in the absence of PDE2A lymphatic junctions fail to stabilize.

**Figure 4:**
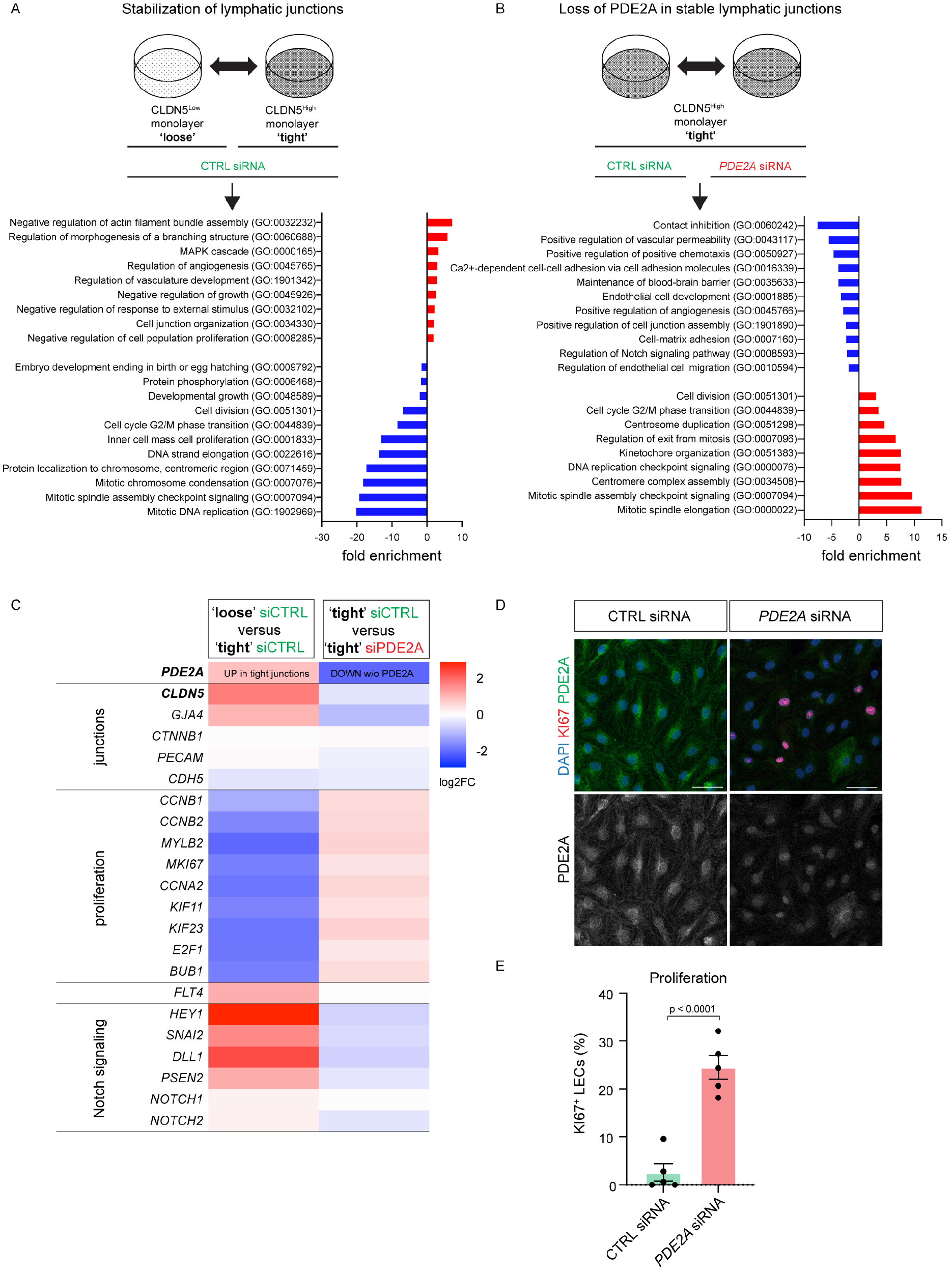
Loss of PDE2A prevents lymphatic contact inhibition. **A)** Global transcriptome analysis comparing HDLECs with “loose” CLDN5^Low^ junctions and “tight” CLDN5^High^ junctions (n = 2 biological replicates) and enrichment of selected GO terms. Up and downregulated GO terms are represented in red and blue, respectively. **B)** Global transcriptome analysis of CTRL and *PDE2A* siRNA treated HDLECs with “tight” CLDN5^High^ junctions (n = 2 biological replicates) and enrichment of selected GO terms. Up and downregulated GO terms are represented in red and blue, respectively. **C)** Selected TOP upregulated and downregulated transcripts regulated by PDE2A are shown. **D)** Representative immunofluorescence staining of 72h CTRL and *PDE2A* siRNA treated HDLECs stained with antibodies against KI67 (magenta), PDE2A (green) and DAPI (blue). Single channel images for PDE2A are shown. Scale bar: 50 μm. **E)** Quantification of the percentage of KI67^+^ HDLECs treated with CTRL and *PDE2A* siRNA (n = 5 independent experiments). Mean ± s.e.m., p-value: Unpaired Student’s *t*-test.

### Loss of PDE2A expression prevents lymphatic EC contact inhibition and cell cycle arrest in vitro

A hallmark of EC junctional maturation is the downregulation of proliferation, contact inhibition, and cell cycle arrest. We therefore sought to investigate if PDE2A might play a role in controlling these processes in LECs. Using an RNAseq approach, we first identified differentially expressed genes potentially involved in junctional maturation and contact inhibition by comparing CTRL siRNA-treated HDLECs with ‘loose’ CLDN5^Low^ junctions and CTRL siRNA-treated HDLECs with ‘tight’ CLDN5^High^ junctions (**Figure 4A**). To identify PDE2A-regulated genes that potentially involved in lymphatic junctional maturation, CTRL siRNA-treated HDLECs with ‘tight’ CLDN5^High^ junctions were further compared to *PDE2A* siRNA-treated HDLECs with ‘tight’ CLDN5^High^ junctions (**Figure 4B**). GO term analysis revealed down-regulation of various GO terms associated with cell cycle transition, cell division and proliferation and up-regulation of GO terms associated with cell junction organization and growth arrest upon junction maturation (**Figure 4A**). In contrast, loss of PDE2A reversed junctional stabilization- and proliferation-associated GO terms (**Figure 4B**), demonstrating a shift of gene expression towards a “loose” junctional state.

*CLDN5* and *GJA4* expression were down-regulated in the absence of PDE2A, while various proliferation markers were up-regulated (**Figure 4C**). Consistently, *PDE2A* siRNA treatment of HDLECs grown to high density maintained the cells in a KI67^+^ proliferative state (**Figure 4D**). Similarly, 5-bromo-2’-deoxyuridine (BrdU) incorporation was maintained only in high density HDLEC monolayers upon *PDE2A* siRNA treatment (**Supplemental figure 5**).

**Figure 5:**
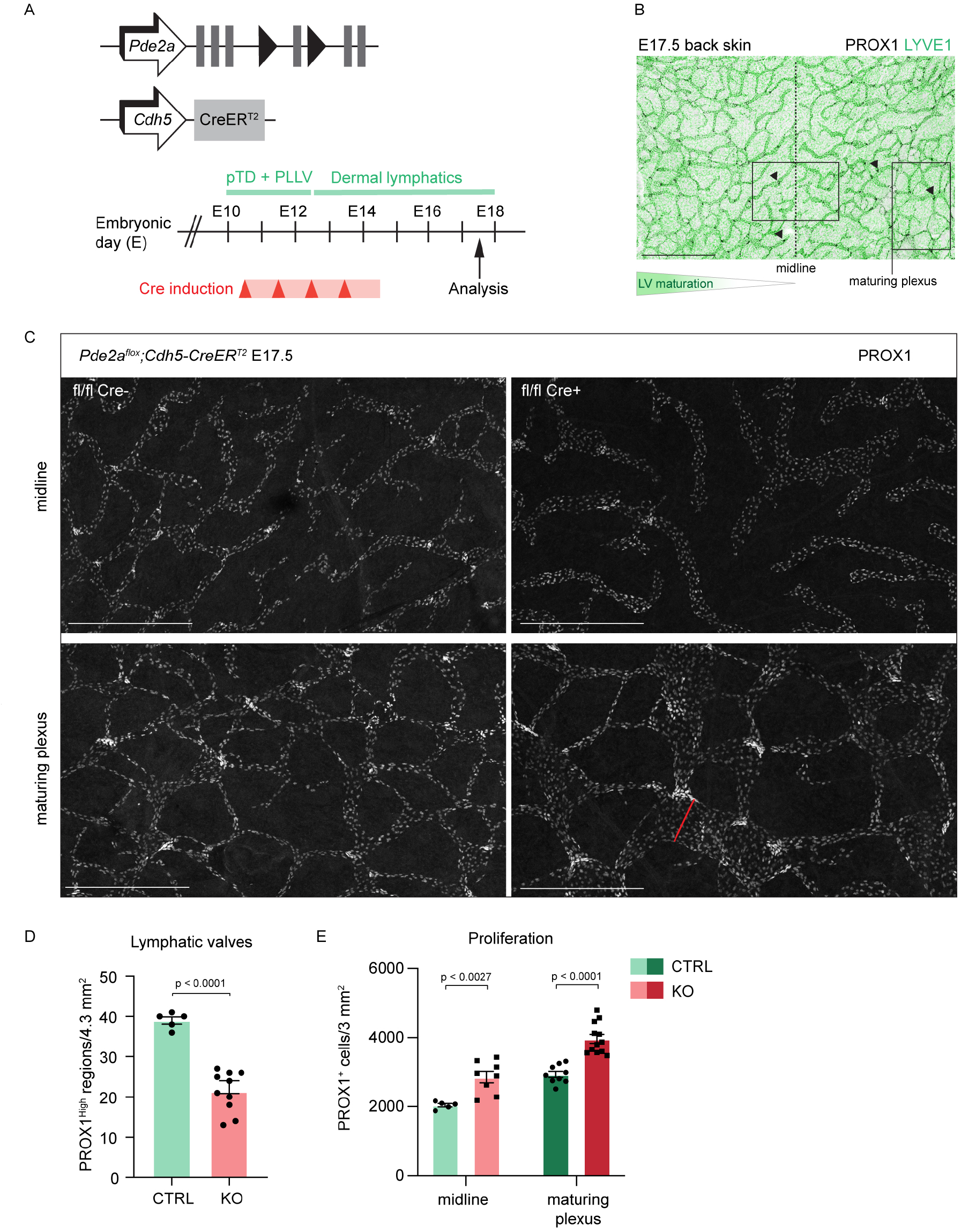
Endothelial-specific *Pde2a* deletion in the *Cdh5*-lineage affects cell cycle arrest and lymphatic maturation at late gestation. **A)** Schematic of the genetic constructs and analyzed embryonic stages. Time points for 4-Hydroxytamoxifen injections (Cre induction) are indicated. **B)** Representative whole-mount immunofluorescence staining of a wild type E17.5 upper thoracic back skin stained with antibodies against PROX1 (grey) and LYVE1 (green) showing the midline region and the maturing lymphatic plexus region analyzed in (C). **C)** Whole-mount immunofluorescence staining of E17.5 upper thoracic back skin midline and maturing plexus from *Pde2a*^*flox/flox*^; *Cdh5-CreER*^*T2*^ (KO) and Pd*e2a*^*flox/flox*^ (CTRL) embryos stained with an antibody against PROX1. Scale bar: 500 μm. **D)** Quantification of lymphatic valves (as PROX1^High^ regions) in the E17.5 upper thoracic back skin of n = 5 KO and n = 3 CTRL embryos. Mean ± s.e.m., p-value: Unpaired Student’s *t*-test. **E**) Quantification of PROX1^+^ cells in midline and maturing plexus of n = 5 KO and n = 3 CTRL embryos. Mean ± s.e.m., p-value: Unpaired Student’s *t*-test.

### Loss of endothelial PDE2A expression prevents lymphatic maturation and contact inhibition at later stages during embryonic development

To avoid early embryonic lethality and analyze lymphatic maturation *in vivo*, we crossed *Pde2a*^*flox*^ mice with tamoxifen-inducible *Cdh5-CreER*^*T2*^ mice. Cre-mediated *Pde2a* deletion was induced from E10.5 for four consecutive days. At that stage non-venous derived LECs had incorporated into the vessels, lateral dermal lymphatics had reached the dorsal midline, valves were formed and lymphatic proliferation was found to decrease in the maturing plexus (**Figure 5A,B**). PROX1 wholemount immunostaining of E17.5 upper thoracic back skins demonstrated reduced number of PROX1^High^ valve regions (**Figure 5C,D**), with increased lymphatic vessel diameter and increased PROX1^+^ cell number in *Pde2a*^*flox/flox*^; *Cdh5-CreER*^*T2*^ mutant embryos compared to Cre-negative control embryos (**Figure 5C,E**). In contrast, sprouting (LYVE1^+^ filopodia) and VEGFR3 expression in the sprouting front and the maturing plexus was not affected even when *Pde2a*^*flox/flox*^; *Cdh5-CreER*^*T2*^ mutant embryos were analyzed at E16.5 before the dorsal midline has closed (**Supplemental figure 6A**). Similarly, *PDE2A* siRNA treatment of HDLECs did not result in changes of VEGFR3 expression (**Figure 4C, Supplemental figure 6B**).

**Figure 6:**
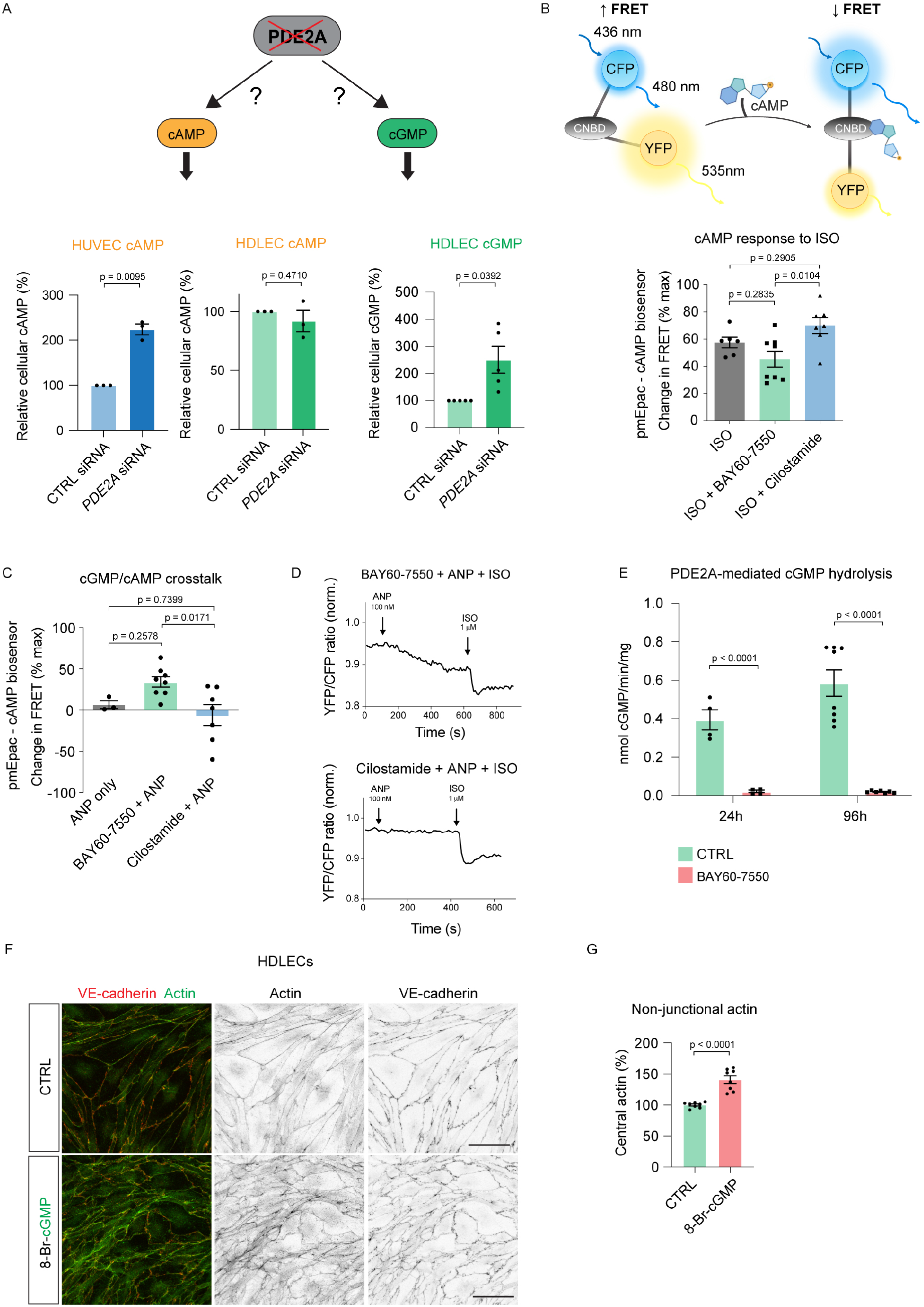
Loss of PDE2A increases lymphatic cGMP levels. **A-left)** Quantification of relative cAMP levels in CTRL and *PDE2A* siRNA treated HDLECs and HUVECs from n = 3 independent experiments per cell type. Mean ± s.e.m., p-value: One-sample t-test. **A-right)** Quantification of relative cGMP levels in CTRL and *PDE2A* siRNA treated HDLECs from n = 5 independent experiments. Mean ± s.e.m., p-value: One-sample *t*-test. **B** Schematic of the FRET Epac1-camps biosensor depicts a conformational change as a result of cAMP binding to the sensor. Quantification of FRET change in HDLECs transduced with the plasma membrane targeted biosensor version called pmEpac1-camps and treated with Isoprenaline (ISO), ISO + BAY60-7550 (PDE2 inhibitor), or ISO + Cilostamide (PDE3 inhibitor). Single points represent single cells from n = 3 independent experiments. Mean ± s.e.m., p-value: One-way ANOVA with Turkey’s multiple comparison test. **C)** Quantification of FRET change in HDLECs transduced with pmEpac1-camps and treated with ANP (atrial natriuretic peptide), BAY60-7550 + ANP, or Cilostamide + ANP. Notably, PDE2 inhibition, but not PDE3 inhibition, significantly increases ANP/cGMP response. Single points represent single cells from n = 3 independent experiments. Mean ± s.e.m., p-value: One-way ANOVA with Turkey’s multiple comparison test. **D)** Representative measurements of FRET change in pmEpac1-camps-transduced HDLECs treated with BAY60-7550 + ANP + ISO or Cilostamide + ANP + ISO. **E)** PDE2A activity measured as a rate of cGMP hydrolysis in HDLECs culture for 24 h or 96 h. Measurements were performed with CTRL or PDE2A inhibition (BAY60-7550). n = 2 independent experiments after 24 h, n = 4 independent experiments after 96 h. Mean ± s.e.m., p-value: Unpaired Student’s *t*-test with Welch’s correction. **F)** Representative immunofluorescence staining of CTRL and 8-Br-cGMP-treated HDLECs stained with an antibody against VE-cadherin (red) and phalloidin (green, to visualize Actin). Single channel images are shown. Scale bar: 50 μm. **G)** Quantification of relative central actin (total actin – junctional actin) from CTRL and 8-Br-cGMP-treated HDLECs from n = 3 independent experiments. Mean ± s.e.m., p-value: Unpaired Student’s *t*-test.

Taken together, in the absence of PDE2A LECs fail to induce contact inhibition and downregulation of proliferation *in vitro* in maturing areas of the embryonic dermal lymphatic network, while sprouting of dermal lymphatics was not affected by PDE2A.

### Loss of PDE2A selectively increases lymphatic cGMP levels in a cAMP-independent manner

Most studies on cAMP function in BECs have shown that increasing exogenous and endogenous cAMP levels stabilize BEC junctions (Beese et al., 2010, Waschke et al., 2004, Fu et al., 2015), while others have reported that prolonged elevation of cAMP destabilizes EC junctions and increases endothelial permeability (Perrot et al., 2018). Consistent with previous data in BECs (Breslin, 2011, Price et al., 2008) we showed that incubation of LECs with cAMP derivatives stabilizes LEC barrier function (**Supplemental figure 7**).

**Figure 7:**
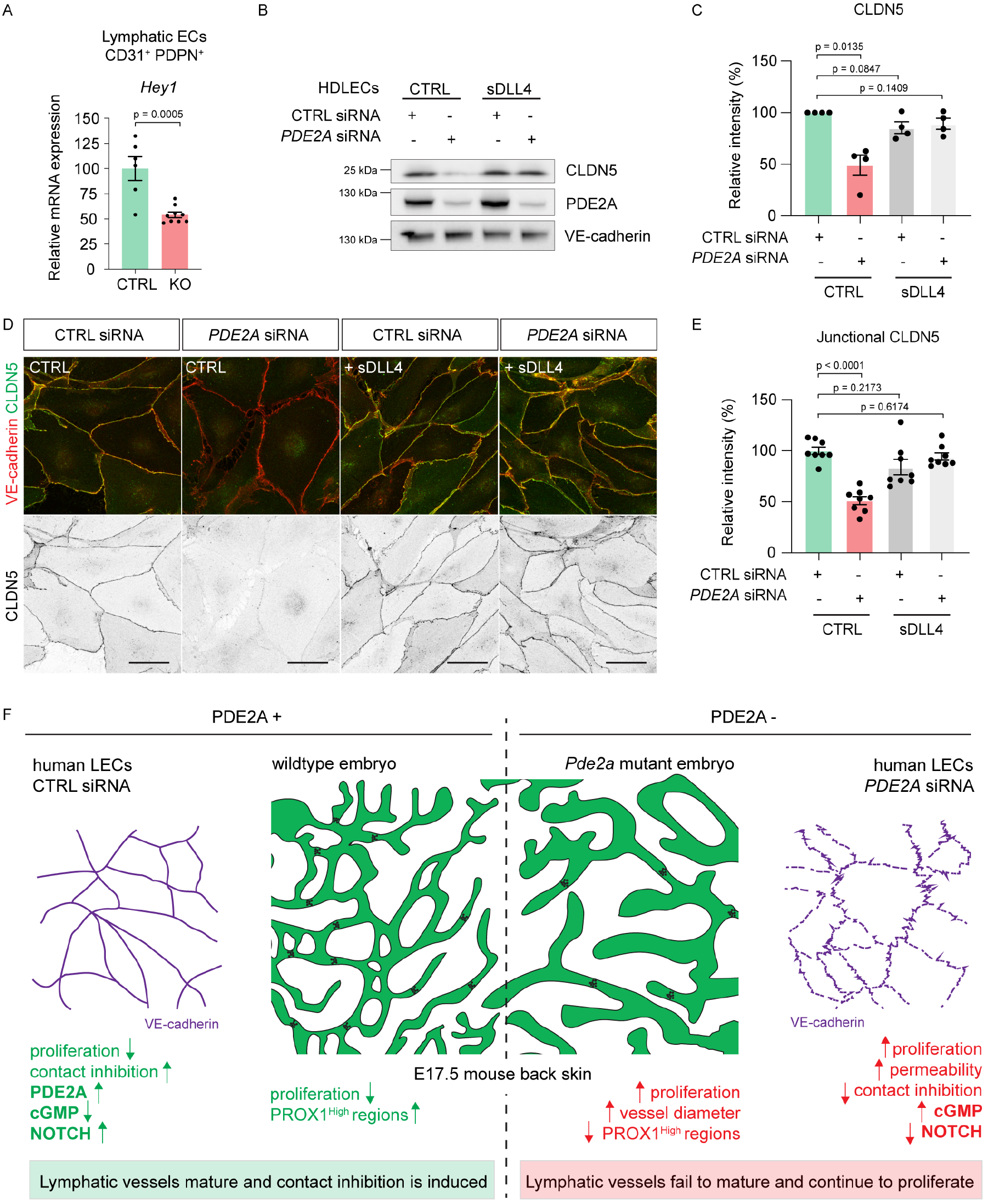
NOTCH signaling activation restores lymphatic junctional maturation. **A)** Relative mRNA expression of the NOTCH target gene *Hey1* in LECs sorted from E14.5 back skins of n = 6 *Pde2a*^*flox/flox*^; *Tie2-Cre* embryos (KO) and n = 9 *Pde2a*^*flox/flox*^ littermate embryos (CTRL). Mean ± s.e.m., p-value: Unpaired Student’s *t*-test. **D**) Western Blot analysis of CTRL and *PDE2A* siRNA treated HDLECs incubated for 24h with or without soluble DLL4 (sDLL4). Lysates were probed with antibodies against CLDN5, PDE2A and VE-cadherin. **E)** Quantification of CLDN5 protein expression from n = 4 independent experiments. Mean ± s.e.m., p-value: One-sample *t*-test. **F)** Immunofluorescence staining of CTRL and *PDE2A* siRNA-treated HDLECs incubated for 24h with or without soluble DLL4 (sDLL4) using antibodies against CLDN5 (green) and VE-cadherin (red). Single channel images are shown for CLDN5. **G)** Quantification of CLDN5 protein in VE-cadherin^+^ junctions from n = 2 independent experiments. Mean ± s.e.m., p-value: One-sample *t*-test. **H)** Schematic of PDE2A-mediated control of lymphatic vessel contact inhibition and maturation.

PDE2A regulates cellular cyclic monophosphate metabolism by catalyzing the hydrolysis of cAMP to AMP and cGMP to GMP, respectively (Zaccolo and Movsesian, 2007, Conti and Beavo, 2007). Loss of PDE2A-mediated catalytic activity is therefore expected to increase lymphatic cAMP and/or cGMP levels that in turn cause lymphatic dysfunction. We therefore sought to analyze if PDE2A differentially regulates cAMP and cGMP levels in HUVECs versus HDLECs. In HUVECs loss of PDE2A function resulted in increased cAMP levels (**Figure 6A**, (Chen et al., 2016)). In contrast, in HDLECs were not altered upon *PDE2A* knockdown (**Figure 6A**). These findings suggested that PDE2A likely regulates lymphatic maturation via the modulation of cGMP levels instead. In agreement with this, siRNA-mediated deletion of *PDE2A* confirmed elevated cGMP levels in HDLECs in comparison to control HDLECs (**Figure 6A**).

In parallel, we used live cell imaging and a plasma membrane targeted biosensor pmEpac1-camps (**Figure 6B**) to measure local cAMP responses at the membrane of HDLECs as previously described for HUVECs (Chen et al., 2016). First, we compared cAMP responses to the beta-adrenergic agonist Isoprenaline (ISO, 1 µM) with and without PDE2 inhibition by BAY60-7550 (100 nM) or PDE3 inhibition by Cilostamide (10 µM). Both inhibitors had no effect on ISO-induced cAMP, as compared to ISO alone (**Figure 6B**). Next, we assessed the effects of PDE2 and PDE3 inhibition on ANP-induced cGMP responses measured as cGMP/cAMP cross-talk by the pmEpac1-camps biosensor (**Figure 6C**). In contrast to intact HUVECs, where PDE3 was the main regulator of membrane cGMP/cAMP crosstalk while PDE2 expression and contribution were negligible (Chen et al., 2016), our measurements in HDLECs showed that PDE2 but not PDE3 inhibition significantly augmented ANP/cGMP responses (**Figure 6C,D**).

Interestingly, we further found that PDE2A inhibition by BAY60-7550 abolished all cGMP-hydrolyzing capacity of LECs at 24h and 96h of culture (**Figure 6E**), demonstrating that under physiological conditions PDE2A plays a central role in the control of cGMP second messenger signaling in LECs. Furthermore, we found that other PDEs, namely PDE12, PDE10A, PDE4B, PDE4D, PDE6D and PDE8A, although being expressed in HDLECs, were not altered upon siRNA-mediated knockdown of *PDE2A* (**Supplemental figure 8A**), suggesting that they are unlikely to compensate for the loss of PDE2A in LECs.

To study if an increase of cGMP in LECs would show junctional disruption similar to the loss of PDE2A, we incubated mature HDLEC monolayers with 250µM 8-Br-cGMP for 48h. As expected, immunofluorescence analysis of VE-cadherin and actin showed disruption of stable HDLEC monolayers with loss of cortical actin upon treatment with 8-Br-cGMP (**Figure 6F,G**).

Conclusively, loss of PDE2A selectively increases lymphatic cGMP but not cAMP levels. Increased LEC cGMP levels result in junctional disruption and a decrease in cortical actin similar to what has been observed upon *PDE2A* deletion.

### PDE2A modulates a crosstalk between cGMP and NOTCH signaling in lymphatic endothelial cells

Last, we sought to analyze how elevated cGMP levels lead to junctional disruption and increased proliferation. GO term analysis highlighted that NOTCH signaling (GO:0008593) was down-regulated upon deletion of *PDE2A* in stable lymphatic junctions (**Figure 4B**). Furthermore, we have previously shown that siRNA-mediated knockdown of *NOTCH1* and NOTCH inhibition via DAPT resulted in increased proliferation (Zhang et al., 2018), suggesting that loss of PDE2A and increased cGMP could regulate lymphatic junctions by interference with NOTCH activation and consecutively NOTCH-induced cell cycle arrest.

Previously, PDE2A and NOTCH signaling have been linked through the miRNA *miR-139-5p* which is located within the second intron of the *Pde2a/PDE2A* gene and regulates NOTCH (Watanabe et al., 2015). However, we did not observe differences in *miR-139-5p* levels in isolated lung ECs of control and mutant *Pde2a*^*flox*^*;Cdh5-CreER*^*T2*^ mice or human LECs treated with CTRL and *PDE2A* siRNA (**Supplemental figure 8B-D**).

In agreement with a potential PDE2A/NOTCH-signaling axis, we found that mRNA expression of the Notch target gene *Hey1* was decreased in sorted CD31^+^; PDPN+ LECs from *Pde2a*^*flox/flox*^; *Tie2-Cre* mutant embryos in comparison to *Pde2a*^*flox/flox*^ control embryos (**Figure 7A**). To demonstrate a functional link of PDE2A and NOTCH signaling in LECs, we analyzed NOTCH signaling in HDLECs upon solubleDLL4 (sDLL4) stimulation dependent on siRNA-mediated *PDE2A* knock-down (**Figure 7B-E**). sDLL4-mediated activation of NOTCH was sufficient to rescue high CLDN5 expression in *PDE2A-*deleted HDLECs suggesting mature lymphatic junctions and intact cell cycle arrest (**Figure 7B,C**). Indeed, immunofluorescence analysis of VE-cadherin and CLDN5 further showed normal junctions in *PDE2A-*deleted HDLECs upon sDLL4-mediated NOTCH activation (**Figure 7D,E**).

Taken together, our study uncovers a mechanism by which PDE2A regulates lymphatic junctional maturation through cGMP-mediated control of Notch signaling (**Figure 7F)**.

## Discussion

During lymphatic development, cell junctions mature and vessel integrity is established to form the lymphatic endothelial barrier. Here, we showed that the phosphodiesterase PDE2A selectively regulates lymphatic vessel maturation during embryonic lymphatic development. Constitutive endothelial deletion of *Pde2a*, prior to lymphatic commitment, resulted in lymphatic dysplasia, while morphology of large blood vessel morphology was not altered. During junctional maturation, PDE2A expression was strongly increased in LECs, but not in venous BECs, and was indispensable for proper lymphatic contact inhibition and maturation *in vitro* and *in vivo*. Mechanistically, PDE2A deletion increased cGMP, but not cAMP levels, leading to a decrease of junction-reinforcing NOTCH signaling activity.

Selective barrier control can be achieved through exclusive expression of junction-controlling molecules in different EC types (Hilfenhaus et al., 2018, Richards et al., 2022). However, other molecules, such as VE-cadherin (Frye et al., 2015, Hägerling et al., 2018) and EphB4 (Frye et al., 2020, Luxán et al., 2019), have been shown to have multifaceted roles in junctional regulation, despite similar expression patterns. Intriguingly, we found that loss of junctional maturation in PDE2A-deficient LECs is caused through accumulation of intracellular cGMP, but not cAMP. This is in line with most studies showing that increased cAMP levels strengthen junctional stability in LECs and BECs (Beese et al., 2010, Waschke et al., 2004, Fu et al., 2015). Similar to PDE2 inhibition (Favot et al., 2003), we show that deletion of PDE2A in HUVECs led to increase cAMP levels, suggesting that in the absence of PDE2A BECs might show altered junctions. Although blood capillary branching was slightly decreased in embryonic back skin upon endothelial *Pde2a* deletion at E14.5, we did not find morphological evidence that BEC junctions or vessel architecture were altered, suggesting that alterations of cAMP levels in BECs might be normalized through other PDEs or endogenous cAMP accumulation is not sufficient to induce BEC dysfunction at this developmental stage. In contrast, compensation of dysregulated lymphatic cGMP levels is not likely to occur because we show for the first time that PDE2A is the only enzyme to hydrolyze cGMP in LECs.

Although we found that loss of PDE2A function does not alter the architecture of blood vessels during embryonic development, PDE2A expression has been shown to be induced under inflammatory conditions in cultured BECs (Surapisitchat et al., 2007, Rentsendorj et al., 2011). In a mouse model of lipopolysaccharide (LPS) treatment and ventilator-induced lung injury (VILI), inhibition of inflammation-induced PDE2A, using an adenovirus expressing a short hairpin RNA, correlated with increased lung cAMP and attenuated disease severity (Rentsendorj et al., 2011). Furthermore, in agreement with our data, inhibition of PDE3A, using Cilostamide, did not alter lymphatic vessel permeability in adult wildtype mice (Scallan et al., 2015). However, inhibition of PDE3A restored mesenteric collecting lymphatic vessel integrity in diabetic mice (Scallan et al., 2015) and improved lymphatic vessel count and flow in a lymphedema mouse model (Kimura et al., 2014). The specific cellular populations responsible for these PDE2A and PDE3A-mediated effects remain to be investigated.

Notably, E14.5 *Pde2a*^*flox/flox*^; *Tie2-Cre* mice exhibited not only a similar lymphatic but also other phenotypes, such as a reduced size of the fetal liver and embryonic death at E14.5-15.5, when compared to the global *Pde2a*^*-/-*^ mice (Assenza et al., 2018, Barbagallo et al., 2020). These findings suggests that the loss of *Pde2a* in the Tie2^+^ lineage is causing embryonic defects and lethality. Future studies will delineate contribution of different cell types of the Tie2^+^ lineage.

NOTCH1 has been shown to regulate general lymphatic growth, sprouting, proliferation and valve morphogenesis (Murtomaki et al., 2013, Zhang et al., 2018, Zheng et al., 2011, Niessen et al., 2011, Fatima et al., 2014). Here, we identified a novel pleiotropic role of NOTCH signaling in LEC contact inhibition, which is modulated through a crosstalk with PDE2A-regulated cGMP signaling. It needs to be further explored if this lymphatic specific crosstalk might be regulated via changes in NOTCH signaling strength (Shen et al., 2021).

An intimate relationship between PDE2A and NOTCH signaling is further supported by the finding that the NOTCH-targeting miRNA *miR-139-5p* (Li et al., 2018) is located within the second intron of the *PDE2A/Pde2a* host gene (Watanabe et al., 2015). *miR-139-5p* has been shown to correlate with *PDE2A* expression, suggesting a potential inhibitory function on this gene (Watanabe et al., 2015). Although we did not expect the second intron of the *Pde2a* host gene to be dysregulated, given that Cre-induced recombination targets exon 4, we analyzed the expression of *miR-139-5p* in isolated lung ECs from 8-week-old control and mutant *Pde2a*^*flox*^; *Cdh5-CreER*^*T2*^ mice. *miR-139-5p* expression was not altered in e*Pde2a* mutant ECs. Additionally, we investigated *miR-139-5p* expression upon siRNA-mediated *PDE2A* deletion in HDLECs to exclude negative feedback on *miR-139-5p* expression. As expected, *miR-139-5p* expression was not altered in CTRL versus *PDE2A* siRNA-treated HDLECs.

Although our assessment of lymphatic PDE2A function was limited to developmental stages, the identified mechanism may be targetable to control lymphatic leakage or regrowth. Altered lymphatic junctional integrity has been implicated in a variety of pathological conditions, highlighting the need to understand the underlying mechanisms. For example, collecting vessel permeability is increased upon infection with *Yersinia pseudotuberculosis* (Fonseca et al., 2015), which results in development of fibrotic processes (Ivanov et al., 2016), increased inflammation and compromised immunity due to reduced flow of lymph and immune cell transport to the lymph nodes (Kuan et al., 2015). Excessive leakage of lymph, caused by dysfunctional lymphatic endothelial junctions in the regions of collecting vessel valves, may also contribute to inherited, primary lymphedema (Mahamud et al., 2019, Sabine et al., 2015). Conversely, controlled disruption of lymphatic contact inhibition via PDE2A inhibition might also contribute to accelerate lymphatic regeneration, e.g., following lymph node dissection.

Up to our knowledge PDE2A has not been implicated in LEC biology. The enzyme is extensively studied in the nervous system, where PDE2A inhibitors have emerged as a novel therapeutic approach to ameliorate cognitive dysfunction in neuropathological disorders, such as schizophrenia or Alzheimer’s disease (Mikami et al., 2017, Baillie et al., 2019, Delhaye and Bardoni, 2021). However, all clinical trials studying PDE2A inhibitors have been unsuccessfully terminated, e.g., due to safety concerns (Baillie et al., 2019, Trabanco et al., 2016). In the light of our findings, PDE2A inhibitors are likely to cause opposing effects dependent on the cell type. Consequently, the identification of ubiquitously expressed molecules with multifaceted roles may also explain and can help to avoid adverse effects of therapeutic drugs targeting.

In summary, this is the first study to identify a role of PDE2A in the lymphatic vasculature. PDE2A controls lymphatic contact inhibition and maturation through NOTCH activity in a cGMP-dependent manner.

## Methods

### Mice

*Tie2-Cre* (B6.Cg-Tg(Tek-cre)1Ywa/J (Kisanuki et al., 2001)), *Cdh5-CreER*^*T2*^ (Tg(Cdh5-cre/ERT2)1Rha (Sörensen et al., 2009)) and *Pde2a* global knockout (B6; 129P2-Pde2A < tm1Dgen>/H; EM: 02366 (Assenza et al., 2018)) mouse lines were previously described. ES cell line containing ‘knockout-first’ *Pde2a* allele (Pde2a^tm1a(EUCOMM)Wtsi^) was obtained from The European Conditional Mouse Mutagenesis Program (EUCOMM). After obtaining germ line transmission, LacZ-neo cassette was removed by crossing with a FLPe deleter strain (Rodríguez et al., 2000) followed by mating to C57BL/6J for at least five generations.

For induction of Cre-mediated recombination in pregnant females, 4-hydroxytamoxifen (4-OHT, Cat#: H7904, Sigma-Aldrich) was dissolved at 10mg/ml in peanut oil and 1mg was administered by intraperitoneal injection from E10.5 for 4 consecutive days. For endothelial cell isolation, we induced Cre recombination in 8-week-old *Pde2a*^*flox*^; *Cdh5-CreER*^*T2*^ mice. Tamoxifen (TAM, Cat#: T5648, Sigma Aldrich) was dissolved at 10mg/ml in peanut oil and 1mg was administered by intraperitoneal injection for 5 consecutive days. ECs were isolated 4 weeks later. All strains were maintained and analyzed on a C57BL/6J background. All the animal procedures conformed to the Directive 2010/63/EU of the European Parliament on the protection of animals used for scientific purposes. Experimental procedures were approved by the Behörde für Justiz und Verbraucherschutz der Freien und Hansestadt Hamburg, Germany (permit number: N070/2019). The animals were kept in the breeding facility of the University Medical Center Hamburg-Eppendorf under animal-friendly husbandry conditions in all phases of life. This applies to the space available, group or individual housing and cage enrichment. Generation of *Pde2a* global knockout and littermate control embryos was conducted with the approval of the Ethic Committee and the Italian Ministry of Health with protocol number n. 919/2020-PR (28/9/2020).

### Cell culture

Primary human dermal lymphatic endothelial cells (HDLECs, Cat#: C-12216) and human umbilical vein endothelial cells (HUVECs, Cat#: C-12203) were obtained from PromoCell. Cells were seeded on tissue culture-treated dishes (Cat#: 430167, Corning^®^) coated with 1 µg/ml bovine fibronectin (Cat#: F1141-1MG, Sigma) diluted in Dulbecco’s phosphate buffered saline (referred to as PBS, Cat#: F1141, R&D Systems).

HDLECs were cultured in complete Endothelial Cell Growth Medium MV2 medium (ECGMV2, Cat#: C-22022, PromoCell) with 25 ng/ml of recombinant human VEGF-C (Cat#: 9199-VC, R&D Systems) and HUVECs were cultured in complete Endothelial Cell Growth Medium (ECGM, Cat#: C-22010, PromoCell). Note, to study PDE2A function *in vitro*, EC monolayer have to be grown to maximum confluency to obtain “*tight junctions*” with high CLDN5 expression.

Murine endothelial cells (mECs) were cultured in mEC medium (0.1% bovine brain extract (BBE)/10% FBS/1%penicillin-streptomycin/1% Non-Essential Amino Acid (NEAA)/Dulbecco Modified Eagle Medium (DMEM) (DMEM Cat#: 41966-029, Gibco; BBE Cat#: CC-4098, Lonza BioScience). All cells were cultured at 37 °C in a humidified atmosphere with 5% CO_2_ (HDLECs and HUVECs) or 10% CO_2_ (mECs).

### Murine endothelial cell isolation

mECs were isolated from lung of *Pde2a*^*flox*^ controls and *Pde2a*^*flox*^; *Cdh5-CreER*^*T2*^ mutants as previously described (Zink et al., 2021). Three lungs isolated from 12-week-old mice were pooled per genotype. Macrovessels were removed and remaining tissue was thoroughly minced using scissors for 15 min. Lung tissue was digested using 1mg/ml Collagenase A (Cat#: 10103586001, Roche; diluted in PBS with 1.25mM CaCl_2_) in a 37°C water bath for 1.5 h, with gentle agitation every 15 min. The cell suspension was then passed through a 40μm cell strainer and single cells were pelleted by centrifugation at 300 x g for 5 min without deceleration. The cell pellet was resuspended in 2mL medium with sheep-anti-rat IgG Dynabeads™ (Cat#: 110-35, Invitrogen) coupled with a CD31 antibody (clone Mec13.3, Cat#: 102501, BioLegend, diluted 1:33). The mix was incubated at 4°C for 45 min with constant rotation. Cells coupled to the CD31-Dynabeads™ were positively selected using a DynaMag™-2 (Cat#: 12321D, Invitrogen) washed twice with mEC medium, resuspended in fresh mEC medium and plated on fibronectin-coated culture dishes, as described above. Lung mECs were sorted again once the monolayer has reached confluency. mECs were then processed for microRNA isolation and Western blot analysis.

### RNA interference

FlexiTube siRNA was used to knock-down human *PDE2A* (Cat#: SI00040159, Qiagen). AllStars Negative CTRL siRNA was used as control (Cat#: 1027281, Qiagen). Cells were subjected to transfection 24h after plating, using Lipofectamine 3000 (Cat#: L3000015, Invitrogen) according to the manufacturer’s instructions.

### Cell growth on stiffness hydrogels

24-well Plate Softwell Easy Coat hydrogels of different stiffness (0.1, 1, 2, 4 and 25 kPa, Cat #SS24-EC, Matrigen) were coated with 10µg/ml collagen I rat tail diluted in PBS (Cat #A10483-01, Gibco) at 4°C and then at 37°C for 1h each. Subsequently, collagen solution was removed and 0.3×10^5^ HDLECs were plated on hydrogels in a volume of 50 µl, allowed to attach at RT for 5 min and then at 37°C for 15 min, before additional culture medium was added. After 24h culture, cells were fixed with 4% PFA in PBS at RT for a total of 20 min, with a change of fixative in between and were then processed for immunofluorescence as described above For RNA sequencing, cells on hydrogels were incubated in OptiMEM Reduced Serum Medium (Cat#:31985-062, Gibco) at 37°C for 30 min. OptiMEM was aspirated, cells were carefully washed with PBS and detached from the hydrogels using 0.05% Trypsin/EDTA (Cat#:25300-054, Gibco). Trypsinization was stopped by adding fresh medium after 5 min, cells were pelleted by centrifugation at 200 x g and RNA isolation was performed as described below.

### BrdU incorporation assay

HDLECs were seeded on 6-well plate dishes at 2×10^5^ cells per well and transfected with CTRL or *PDE2A* siRNA as described above. The day after transfection, cells were trypsinized and seeded in 8-well Lab-Tek II chamber slides (Cat#: 154941, Lab-Tek®) at concentrations of 2×10^4^, 5×10^4^ and 1×10^5^ cells per well. After 36h ECGMV2 was changed to basal medium (ECGMV2 without supplements, Cat#: 22221, PromoCell) for 4h. Next, 3µg/ml BrdU (Cat#: 347580, BD) with 100ng/ml VEGF-C (R&D Systems) in basal medium were then added to the cells for 12h. Subsequently, cells were fixed with 4% PFA in PBS for 20 min at RT. Samples were washed once with PBS and incubated with 2N HCL at RT for antigen retrieval for 30 min, followed by immunofluorescence analysis.

### *in vitro* permeability assay

To determine paracellular permeability, 4×10^4^ CTRL or *PDE2A* siRNA treated HDLECs were seeded on fibronectin–coated Transwell filters (Costar 3413, 0.4 μm pore size; Corning) and grown to confluency. When maximum confluency was reached, 0.25mg/ml FITC-dextran (40 kDa, Cat#: FD40S, Sigma-Aldrich) diluted in ECGMV2 was added to the upper chamber of the Transwell filters and monolayer diffusion was allowed for 1h. Fluorescence in the lower chamber was measured with a Tecan Spark® 10M microplate reader, and monolayer integrity was confirmed by immunofluorescence staining for VE-cadherin after each assay.

### cAMP and cGMP incubation

1×10^5^ HDLECs were seeded per well in an 8-well Lab-Tek II chamber slide (Cat#: 154941). After 48h culture, cells were incubated with 250µM 8-Br-cAMP (Cat#: B5386, Sigma-Aldrich) or 250µM 8-Br-cGMP (Cat#: B1381, Sigma-Aldrich) for 1h and 48h, respectively. Subsequently, cells were fixed with 4% PFA in PBS at RT for 20 min. Samples were washed once with PBS followed by immunofluorescence analysis as described above.

### cAMP/cGMP quantification

cAMP and cGMP measurements were performed using Direct cAMP ELISA Kit (Cat#: ADI-900-066, Enzo Life Science, Inc.) and Direct cGMP ELISA Kit (Cat#: ADI-900-014, Enzo Life Science, Inc.). In both cases, the acetylated version of the assays was performed. HDLECs and HUVECs were seeded on fibronectin-coated 6-well plates at 2×10^5^ cells per well and transfected with CTRL or *PDE2A* siRNA. After 72h, cell pellets were collected, snap frozen and stored at -80°C. To measure cAMP/cGMP, cells were lysed in 300μL lysis buffer (0.1N HCL/ 0.1% Triton-X100) at RT for 10 min. After centrifugation at 9,500 x g for 10 minutes, 250μL of samples were transferred to a new tube and processed according to the manufacturer’s instructions. Optical density was measured with a Tecan Spark® 10M microplate reader.

### Phosphodiesterase activity assay

Cells were homogenized in 20 mM Tris-HCl buffer (pH 7.2, 0.2 mM EGTA, 5 mM β-mercaptoethanol, 2% (v/v) antiprotease cocktail (Merck KGaA, Germany), 1 mM PMSF, 5 mM MgCl2, 0.1 % (v/v) Triton X-100 and centrifuged at 14,000 × g for 30 min at 4 °C. PDE activity was measured on the supernatant according to the method described by Thompson and Appleman (Thompson and Appleman, 1971) in 60 mM Hepes (pH 7.2, 0.1 mM EGTA, 5 mM MgCl2, 0.5 mg/ml bovine serum albumin and 30 mg/ml soybean trypsin inhibitor), in a final volume of 0.15 ml. The reaction was started by adding tritiated substrate ([3H] cGMP) at a final concentration of 1 μM. The reaction was stopped by adding 50 μl of 0.1 N HCl and then neutralized with 50 μl of 0.1 N NaOH in 0.1 M Tris-HCl pH 8.0. Subsequently, 25 μl of 2 mg/ml of 5ʹ-nucleotidase (snake venom from Crotalus atrox; Merck KGaA) in 0.1 M Tris-HCl pH 8.0 were added. Samples were gently mixed and incubated at 30°C for 30 min to allow complete conversion of 5’-nucleotide to its corresponding nucleoside. Unhydrolyzed cyclic nucleotide and the corresponding nucleoside were separated by DEAE-Sephadex A-25 columns. The eluate was mixed with ULTIMA GOLD scintillation liquid (PerkinElmer, USA) and counted on a Tri-Carb 2100TR Liquid Scintillation Counter (2000CA; Packard Instruments, USA). To evaluate the enzymatic activity of PDE2A, the specific inhibitor BAY 60-7550 was added to the reaction mix at a final concentration of 0.1 μM.

### Notch activation assay

To activate Notch signaling in LECs, 2×10^5^ HDLECs per well were seeded in 6-well plate dishes, transfected with CTRL or *PDE2A* siRNA and, after 72h, treated twice every 12 h with human recombinant sDLL4 (1µg/ml, Cat#: SRP3026, Sigma Aldrich) for a total of 24h, followed by protein isolation.

### RNA sequencing

RNA sequencing (RNAseq) was performed by BGI, Shenzhen, China. Briefly, mRNA molecules were purified from total RNA using oligo(dT)-attached magnetic beads and fragmented into small pieces using mechanical fragmentation (6min with 85℃). First-strand cDNA was generated using random hexamer-primed reverse transcription, followed by a second-strand cDNA synthesis. The synthesized cDNA was subjected to end-repair and was 3’-adenylated. Adapters were ligated to the ends of the 3’-adenylated cDNA fragments. The adapters were used to amplify cDNA fragments using PCR. PCR products were purified with Ampure XP Beads (Agencourt, Beckman Coulter, Krefeld, Germany), and dissolved in EB buffer. The library was validated on the Agilent Technologies 2100 bioanalyzer (Agilent, Santa Clara, US). Next, the double stranded PCR products were heat-denatured and circularized by the splint oligo sequence. The single strand circle DNA (ssCirDNA) were formatted as the final library. For sequencing, the library was amplified with phi29 DNA polymerase to generate DNA nanoballs (DNB) which had more than 300 copies of one DNA fragment. The DNBs were loaded into the patterned nanoarray and pair end 100 bases reads were generated using sequencing by synthesis.

The quality of the sequence reads was assessed by FastQC (v0.11.9). Trimmomatic (v0.36) was employed to remove sequencing adapters and low-quality bases (Phred quality score below 20) from the 3’-end of the sequence reads (Bolger et al., 2014). Thereafter, reads were aligned to the human reference assembly (GRCh38.98) using STAR (v2.7.3a) (Dobin et al., 2013). Batch effect correction was applied with ComBat-seq (Zhang et al., 2020b). Differential expression was assessed with DESeq2 (Love et al., 2014), a gene was considered differentially expressed if the corresponding False Discovery Rate (FDR) was lower than or equal to 0.1 and the absolute log2-transformed fold change (log2FC) was higher than or equal to 0.5. The detection of overrepresented pathways was performed using WebGestalt platform 2019 (Liao et al., 2019).

### Immunofluorescence

For immunostaining of whole-mount back skins and 100μm vibratome sections, tissues were fixed in 4% PFA overnight at 4°C, permeabilized in 0.5% Triton-X100/PBS and blocked in 0.3% Triton-X100/PBS (PBSTx)/3% BSA (blocking buffer). Primary antibodies were incubated in blocking buffer at 4°C overnight. After washing in PBSTx, the samples were incubated with Alexa Fluor (AF)-conjugated secondary antibodies in blocking buffer at 4°C overnight. After further washing, the tissue samples were mounted in Fluoroshield Mounting Medium without DAPI (Cat#: F6182, Sigma-Aldrich).

For staining of HDLECs and HUVECs, cells were seeded in 8-well Lab-Tek II chamber slides (Cat#: 154941, Lab-Tek^®^) at the desired concentration (from 2×10^4^ to 1×10^5^ cells per well) and transfected with CTRL or *PDE2A* siRNA as described above. 72h post-transfection, cells were fixed with 4% PFA in PBS at RT for 20 min or with ice-cold methanol (for CLDN5 immunostaining) at 4°C for 20 min. Samples were permeabilized using 0.5% Triton-X100/PBS for 5 min followed by blocking with 2% BSA/PBSTx for 1h. Primary antibodies were incubated for 1h, followed by washing twice with PBSTx and subsequently cells were incubated with secondary antibodies for 45 min before further washing and mounting in Fluoroshield Mounting Medium with DAPI (Cat#: F6057, Sigma-Aldrich). Staining was performed at RT if not specified otherwise.

The following antibodies were used: goat anti-PROX1 (Cat#: AF2727, R&D Systems, 1:100), rat anti-Endomucin (Cat#: sc-65495, Santa Cruz Biotechnologies, 1:100), rat anti-CD31 (Cat#: 102501, BioLegend, 1:100), mouse anti-PDE2A (Cat#: sc-271394, Santa Cruz Biotechnologies, 1:50), rabbit anti-KI67 (Cat#: ab16667, abcam, 1:200), goat anti-NRP2 (Cat#: AF567, R&D Systems, 1:20), goat anti-VEGFR3 (Cat#: AF743, R&D Systems, 1:100) mouse anti-VE-cadherin (F8, Cat#: sc-9989, Santa Cruz Biotechnology, 1:100), goat anti-VE-cadherin (Cat#: AF1002, R&D Systems, 1:200) rabbit anti-LYVE1 (Cat#: 103-PA50AG, Relia Tech GmbH, 1:200), rabbit anti-CLDN5 (34–1600, Thermo Fisher Scientific,1:50) and mouse anti-BrdU (Cat#: 347580, BD Biosciences, 1:2.5 in 3% BSA/PBSTx). Secondary antibodies conjugated to AF488, AF594 and AF647 were obtained from Jackson ImmunoResearch (all used 1:200). Additionally, AF594- or AF488 Phalloidin (Cat#: A12381, Cat#: A12379, Thermo Fisher Scientific, 1:40) were used.

### Western blot

Total protein extract was obtained adding 200µl ice-cold pTyr lysis buffer (20mM Tris-HCL pH7.4/ 150mM NaCl/ 2mM CaCl_2_/ 1.5mM MgCl2/ 1% Triton-x-100/ 0.04% NaN_3_/ EDTA-free protease inhibitors) to the cells plated on fibronectin-coated 6-well plates. Cell lysates were incubated at 4°C in overhead rotation for 30 min, then centrifuged to discard cell debris at 14,000 x g, 4°C, for 15 min. Supernatants were collected and protein concentration was determined using a Pierce™ BCA Protein Assay Kit (Cat#: 23225, Thermo Fisher scientific) according to manufacturer’s instructions. Subsequently, equal amounts of proteins were denaturated using 4xLaemmli sample buffer with 10% β-mercaptoethanol at 95°C for 5 min, then separated via SDS-PAGE. Proteins were transferred to a Immobilion®-P Transfer Membrane (0.45 µm pore size, Cat#: IPVH00010, Merck) and blocked for 1h at RT in 1xTris Buffered Saline Tween (TBST) (150mM NaCl/ 10mM Tris-HCl (pH 7.4)/ 0.05% Tween/ 5% (w/v) milk powder). The membranes were subjected to overnight incubation at 4°C with primary antibodies diluted in 1xTBST/5% BSA. Membranes were rinsed three times with 1x TBST for 10 min each and incubated with HRP-conjugated secondary antibodies (diluted in TBST/5% (w/v) milk powder) at RT for 1h. Membranes were rinsed three times with TBST for 10 min each and specific binding was detected using the enhanced chemiluminescence (ECL)™ Select Western Blotting Detection Reagent (Cat#: RPN2235, Cytiva) and the Gel Doc™ MP Imaging System (BioRad). Protein molecular masses were estimated relatively to the electrophoretic mobility of co-transferred prestained protein marker PageRuler™ Prestained Protein Ladder (Cat#:26616, Thermo Fisher Scientific). The following primary antibodies were used: mouse anti-PDE2A (Cat#: sc-271394, Santa Cruz Biotechnologies, 1:300), rabbit anti-PDE3A (kind gift of Chen Yan (Bork et al., 2021), 1:1000), goat anti-VE-cadherin (C19 clone, Cat#: sc-6458, Santa Cruz Biotechnologies, 1:250), goat anti-VE-cadherin (Cat#: AF1002, R&D Systems, 1:1000), rabbit anti-CLDN5 (Cat#: 34-1600, Thermo Fisher Scientific, 1:500), rabbit anti-VEGFR3 (Cat#: MAB3757, Millipore, 1:500), rabbit anti-β-actin (Cat#: 4970, Cell Signaling, 1:1000), and anti-PROX1 (Cat#: AF2727, R&D Systems, 1:200).

### Flow cytometry

Back skins were collected from E14.5 *Pde2a; Tie2-Cre* embryos, dissected in ice-cold PBS and then digested with 4mg/ml Collagenase Type IV/ 0.2mg/ml DNase I /10% FCS/PBS under constant shaking with 700 rpm at 37°C for 12-15 min (Collagenase Type IV, Cat#: 17104-019, Gibco; DNase I, Cat#: 10104159001, Roche). Samples were filtered using a 70µm nylon filter and then washed twice with FACS buffer (0.5% FCS/2mM EDTA in PBS) to quench enzymatic activity by dilution, before the cell pellet was resuspended in 100µl Fc block (mouse BD Fc Block CD16/CD32 Clone 2.4G2, Cat#: 553141, BD Pharminogen, 1:100 in FACS buffer) and incubated on ice for 10 min. Next, cells were stained with CD31-PE-Cy7 (clone 390, Cat#: 25-0311-82, Invitrogen eBioscience, 1:300), PDPN-APC (clone 8.1.1, Cat#: 127410, Biolegend, 1:200), CD45-eF450 (Cat#: 48-0451-82, Invitrogen eBioscience, 1:20), Ter119-eF450 (Cat#: 48-5921-82, Invitrogen eBioscience, 1:20) and CD11b-eF450 (Cat#: 48-0112-82, Invitrogen eBioscience, 1:20) on ice for 15 min. Dead cells were labelled using 1µM SytoxBlue (Cat#: S34857, Invitrogen). Cells were re-filtered and directly subjected to sorting with a BD Biosciences FACSAria™IIIu run with Diva 8.0.1 software. Single cells were gated from FSC-A/FSC-H plots, followed by exclusion of dead cells and immune cells (CD45^+^/Ter119^+^/CD11b^+^). ECs were gated based on CD31 and LECs (CD31^+^/PDPN^+^) were collected directly in RLT lysis buffer for RNA isolation. Compensation was achieved using UltraComp eBeads Plus Compensation Beads (Cat# 01-3333-42, Invitrogen). If not stated otherwise samples were incubated and handled on ice.

### qRT-PCR analysis

For qRT-PCR analysis of HDLECs, HUVECs and mECs, total RNA was isolated by RNeasy® Mini Kit (Cat#: 74104, Qiagen). In case of sorted murine ECs RNAeasy Micro Kit (Cat#: 74004, Qiagen) was used. RNA was reverse transcribed using SuperScript® IV VILO cDNA Synthesis Kit (Cat#: 11754050, Thermo Fisher Scientific) according to the manufacturer’s instructions. cDNA from sorted murine ECs was additionally pre-amplified using the TaqMan PreAmp Master Mix (Cat#: 4391128, Applied Biosystems). Gene expression levels were analyzed using TaqMan Gene Expression Assay (Cat#: 4369016, Applied Biosystems) and a StepOnePlus RT PCR system (Applied Biosystems). Relative gene expression levels were normalized to *GAPDH*. The following probes were used for human samples (all from Thermo Fischer Scientific): Hs99999905 *GAPDH*, Hs00159935_m1 *PDE2A*, Hs01012698_m1 *PDE3A*, Hs01098928_m1 *PDE10A*, Hs00698272_m1 *PDE12*, Hs00963643_m1 *PDE4B*, Hs01579625_m1 *PDE4D*, Hs01062025_m1 *PDE6D* and Hs01079617_m1 *PDE8A*. For murine cells, the following probes were used (all from Thermo Fischer Scientific): Mm99999915_g1 *Gapdh*, Mm01136644_m1 *Pde2a* and Mm00468865_m1 *Hey*.

### microRNA expression analysis

HDLECs were seeded in 6-well plate dishes with 2×10^5^ cells per well and transfected with CTRL or *PDE2A* siRNA as described above. At 72h post-transfection, microRNA (miRNA) was isolated using miRNeasy Mini Kit (Cat#: 217004, Qiagen) and reverse transcribed using TaqMan MicroRNA Reverse Transcription Kit (Cat#: 4366596, Thermo Fisher Scientific) according to the manufacturer’s instructions. miRNA was isolated from lung mECs of *Pde2a*^*flox*^ controls and *Pde2a*^*flox*^; *Cdh5-CreER*^*T2*^ mutants and processed similarly.

*miR-139-5p* expression levels were analyzed using TaqMan MicroRNA Assay (Cat#: 4427975, Thermo Fisher Scientific) and a TaqMan Universal Master Mix II no UNG with a StepOnePlus RT PCR system (Applied Biosystems). Relative microRNA expression levels were normalized to *RNU48* (assay ID: 001006) for HDLECs, or *snoRNA202* (assay ID: 001232) and *snoRNA234* (assay ID: 001234) for mECs (all from Thermo Fisher Scientific) (Wong et al., 2007). The following probe was used for both sample types: MI0000693 *miR-139-5p*.

### Förster resonance energy transfer (FRET) measurements

HDLECs were seeded on 25mm fibronectin-coated glass coverslips. Following 3 days in culture, cells were transduced with an adenoviral vector to express FRET-based biosensor pmEpac1-camps, targeted the plasma membrane (Perera et al., 2015), with multiplicity infection of 0.1. After 40 to 48h, cells with sufficient sensor expression were used for FRET measurements performed on inverted fluorescent microscope Leica DMI 3000 B with oil-immersion 63x/1.40 objective and the MicroManager 1.4 software. FRET setup included CoolLED at 440nm, beam-splitter DV2 Dual View (Photometrics), and CMOS (OptiMOS, QImaging) camera chip to record images every 10 seconds. Tested compounds (BAY60-7550; Cat# sc396772, Santa Cruz, Cilostamide; Cat# sc201180A, Santa Cruz, human Atrial natriuretic peptide (ANP; Cat# 4011941, Bachem), Isoprenaline; Cat# I6504, Sigma Aldrich) were diluted in FRET buffer (144 mM NaCl, 5.4 mM KCl, 1 mM, MgCl_2_, 1 mM CaCl_2_, 10 mM HEPES; pH 7.3). ImageJ was used for the analysis of raw data which were corrected offline in Microsoft Excel as previously described (Börner et al., 2011).

### Image acquisition

Confocal images of whole-mount back skins, vibratome sections and *in vitro* experiments represent maximum intensity projections of Z-stacks that were acquired using a Leica SP8 inverted microscope with HCX PL APO CS 10x/0.40 DRY or HC PL APO CS2 63x/1.30 GLYC objectives and Leica LAS-X software. Stereomicroscope images of embryos were acquired with Zeiss Stemi 508 with Axiocam 208 Color. Brightfield images of spheroids were taken using Nikon Eclipse Ts2R microscope run with NIS Elements BR Software (Version 5.11, Nikon).

### Image quantification

All quantifications were done using Fiji ImageJ unless stated otherwise. For quantification of central actin, a threshold for junctional VE-cadherin staining was applied, a threshold-based mask was created and pixel intensities of the junctional actin fraction and overall actin pixel intensity was subtracted. 4–11 images (250µm^2^, maximum intensity projection images with 12 z-stacks) were acquired from n=2 independent experiments.

For junctional CLDN5 immunostaining pixel intensity measurements, a defined threshold for junctional VE-cadherin staining was applied, a threshold-based mask was created and pixel intensities of CLDN5 immunostaining in the junctions was analyzed. CLDN5 pixel intensities (integrated density) were detected from 6 images from n = 2 independent experiments. Quantification of proliferating HDLECs was performed by manual counting of KI67+ or BrdU+ cells over the total number of cells per field of view using DAPI signal. 4-5 regions for each condition were imaged and quantified from n=5 independent experiments for KI67 staining, n=3 independent experiments for BrdU assay at 2×10^4^ and 1×10^5^ cells per well and n=2 independent experiments for BrdU assay at 5×10^4^ cells per well.

For quantification of the number and length of LEC sprouts in the fibrin gel spheroid assay, phase contrast images were used. Cumulative sprout length was determined by adding the lengths of each spheroid. 20–30 spheroids from n=2 independent experiments were analyzed for each condition.

Quantification of lymphatic vessel diameter and avascular midline was done using maximum intensity projection images of tile-scanned E14.5 back skin samples (xy = 4800μm x 1000μm, upper thoracic region). For measurement of the distance of lymphatic sprouts from the midline (i.e. avascular midline), dorsal midline was marked with a line in the upper thoracic region and 2–4 measurements on each side of the midline were taken from n=16 *Pde2a*^*flox*^ control embryos and n=13 *Pde2a*^*flox*^; *Tie2-Cre* mutant embryos. Clusters of LECs not connected to the proximal lymphatic network were not taken into account for measurements. Lymphatic vessel diameter was determined by measuring the thickest part of each vessel segment in between branch points. 20–30 measurements in the upper thoracic region were taken from n=15 *Pde2a*^*flox*^ control embryos and n=9 *Pde2a*^*flox*^; *Tie2-Cre* mutant embryos. For blood capillary branching measurement in the back skins, midline region without lymphatic vessels was considered (xy = 2000μm x 860μm) and analyzed using the Skeletonize plugin in ImageJ. The final number of branches was calculated as branch points/mm^2^ in n=10 *Pde2a*^*flox*^ control embryos and n=8 *Pde2a*^*flox*^; *Tie2-Cre* mutant embryos.

The quantification of jugular lymph sac lumen was performed using the ImageJ area measurement plugin with 1-4 measurements per embryo and n = 4 *Pde2a*^*flox*^ control and n = 4 *Pde2a*^*flox*^; *Tie2-Cre* mutant embryos.

For quantifications of PROX1+ cells in E17.5 back skins, 2-3 z-stack images (xy = 2000μm x 1500μm) per region (midline and maturing plexus) were analyzed from n=3 *Pde2a*^*flox*^ control embryos and n=5 *Pde2a*^*flox*^; *Cdh5-CreER*^*T2*^ mutant embryos. A threshold for nuclear PROX1 staining was applied and particle analysis was performed to count positive nuclei. For quantifications of PROX1^High^ valve regions in E17.5 back skins, 1-2 regions of a 2-3 z-stack image (xy = 2900μm x 1500μm) were analyzed from n=3 *Pde2a*^*flox*^ control embryos and n=5 *Pde2a*^*flox*^; *Cdh5-CreER*^*T2*^ mutant embryos. Western blot signal quantifications were done using BioRad Image Lab Software.

### RNA single cell sequencing data analysis

The cellular expression of *Pde2a* in mouse lung tissue was extracted from the publicly available processed data and metadata present in the Tabula Muris data set (deposited on Figshare) (Schaum et al., 2018). The data were normalized using the NormalizeData function as implemented in the Seurat package (Satija et al., 2015). The normalized data were autoscaled, and principal component analysis (PCA) was performed on variable genes followed by Uniform Manifold Approximation and Projection (UMAP) visualization (umap package) (McInnes et al., 2018). Clusters were annotated based on literature-curated marker genes of cell phenotypes. The R implementation of the Plotly software (https://plotly.com/r/) was used for violin and dot plot visualization.

### Statistical analysis

GraphPad Prism 9 was used for graphic representation and statistical analysis of the data. We used 2-tailed unpaired Student’s t-test to compare between two means (assuming equal variance) and including Welch’s correction (not assuming equal variance), One-sample t-test to compare sample mean with a normalized control value = 1 or 100, and One-way ANOVA with Turkey’s multiple comparison test for comparison of multiple conditions. Sorted murine LECs and BECs mRNA levels were normalized using GAPDH as housekeeping gene, 2^(-dCT) calculated as a measure of mRNA expression per analyzed sample and presented as the percentage of mRNA relative to the average of the (respective) controls. Differences were considered statistically significant when p<0.05.

## Acknowledgements

We thank Ralf Adams (Max Planck Institute for Molecular Biomedicine, Münster) for the *Cdh5-CreER*^*T2*^ mice. We also thank Richard Dabels, Nicole Lüder, May Cathleen Müller and Krimhild Scheike for technical assistance and the Microscopy Imaging Facility (DFG Research Infrastructure Portal: RI_00489) at the University Medical Center Hamburg for technical microscopy support. This work was supported by the Werner Otto Medical Foundation Hamburg (8/95), by an Exploration Grant of the Boehringer Ingelheim Foundation (BIS) and the German Research Foundation (DFG) grant FR4239/1-1 to M.F.

## Author contributions

Conceptualization, M.F.; Investigation, C.C., L.L., S.A.H., D.S., R.K., S.C. and M.F.; Resources, T.S., M.P., V.N., T.R. and M.F.; Writing – Original Draft, C.C. and M.F.; Writing – Review & Editing, C.C., L.L., S.A.H., D.S., C.S., R.K., S.C., J.K., CH.S., M.B., R.M., T.S., M.P., V.O.N., T.R. and M.F.; Visualization, C.C., L.L., S.A.H., R.K. and M.F.; Formal Analysis, C.S., J.K., CH.S. and M.F.; Supervision, M.P., V.O.N. and M.F.; Project Administration, M.F.; Funding Acquisition, M.F.

## Data and code availability

The authors declare that the data supporting the findings of this study are available within the paper and its supplementary information. Bulk RNA sequencing data reported in this publication have been submitted to the European Nucleotide Archive (ENA). They are publicly available under accession PRJEB57972. Tabula muris data were downloaded from NCBI GEO database (accession number: GSE109774). Further information and requests for resources and reagents should be directed to and will be fulfilled by Maike Frye.

