## Supplemental information for "The phosphodiesterase 2A regulates lymphatic endothelial development via cGMP-mediated control of Notch signaling"

Carlantoni et al.

Supplemental information

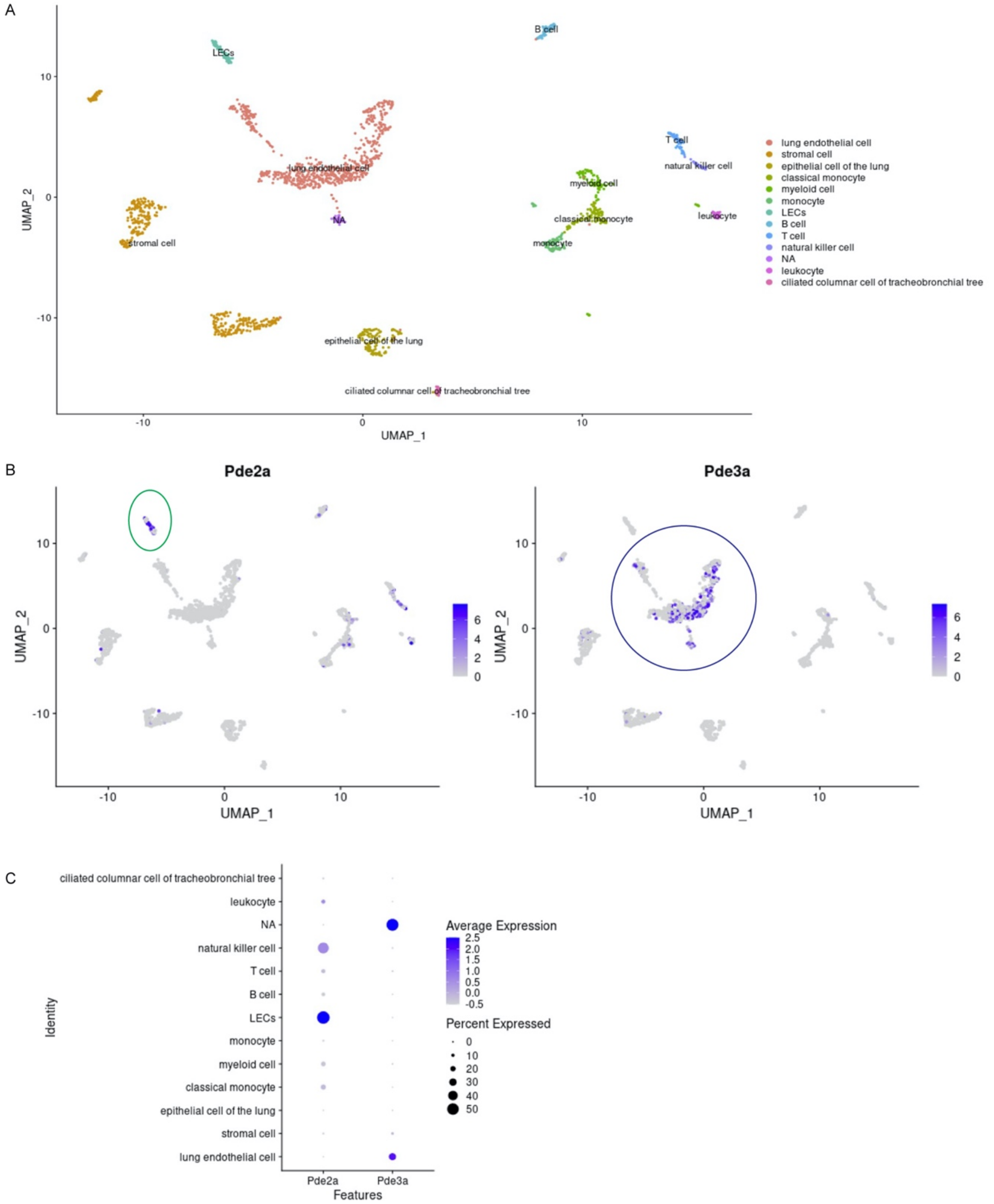


**Supplemental figure 1: *Pde2a* is enriched in lymphatic endothelial cells in adult mice.**
**A**) Uniform manifold approximation and projection (UMAP) representation of the lung cell of Tabula Muris datasets. Each dot represents a single cell. Based on the gene expression, cells were labeled as discrete cell types. **B**) UMAP plot, showing expression of *Pde2a* (left) and *Pde3a* (right) in lung cells. Color-coding indicates expression of representative marker genes for each cell type. Color scale: purple, high expression; grey, low expression. **C**) Dot-plot heatmap of *Pde2a* (left) and *Pde3a* (right). The color intensity of each dot represents the average level of marker expression, whereas the dot size reflects the percentage of cells expressing the marker within given cluster. LECs, lymphatic endothelial cells.

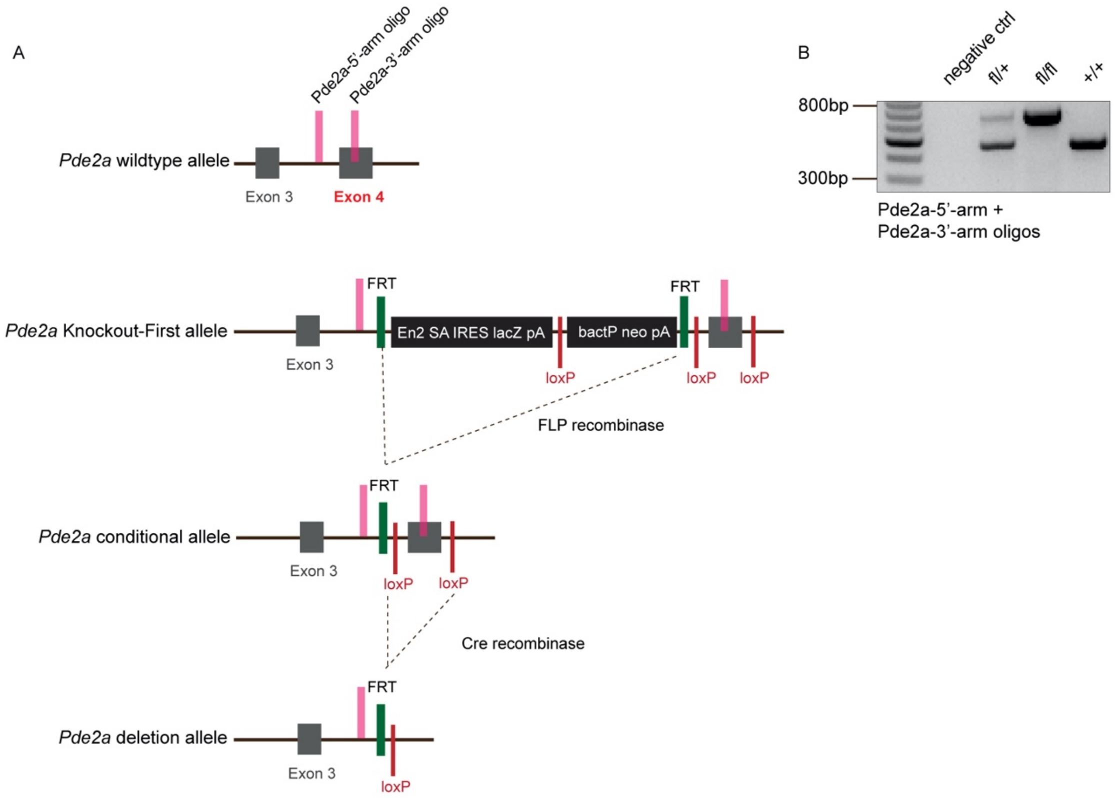


**Supplemental figure 2: Generation of conditional *Pde2a* knockout mice.**
**A**) Schematic of the *Pde2a* wild type allele, targeted 'Knockout-First' allele, conditional allele (floxed) and deletion allele. The 'Knockout-First' allele ($Pde2a^{tm1a(EUCOMM)Wtsi}$) contains an IRES:lacZ trapping cassette and a floxed promoter-driven neo cassette. Flp converts the 'Knockout-First' allele to a conditional allele. Cre deletes the floxed exon 4, generating a null allele and disrupting *Pde2a* gene function. The locations of PCR genotyping primers are indicated in pink. **B**) PCR genotyping of wild type and conditional floxed *Pde2a* mice.

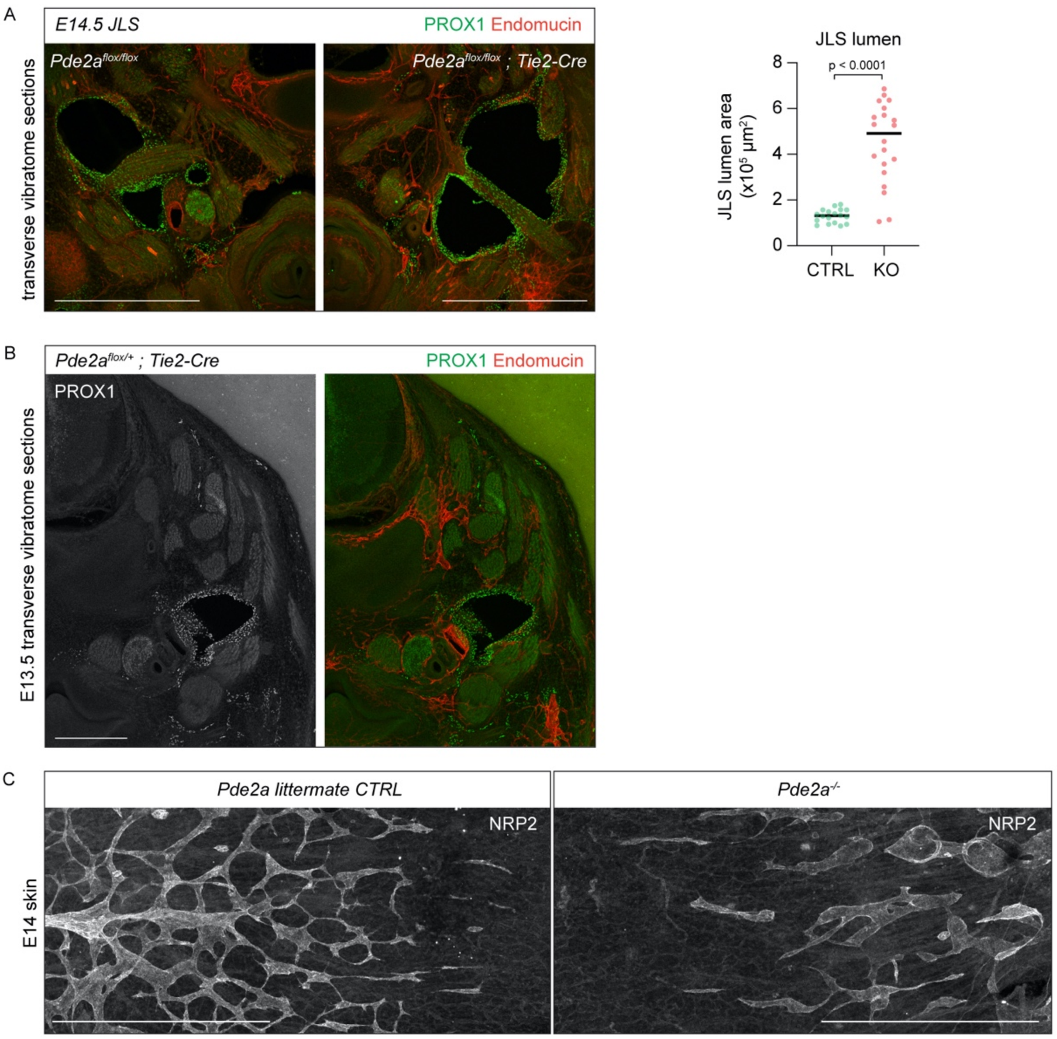


**Supplemental figure 3: Endothelial *Pde2a* deletion results in dysregulation of embryonic lymphatic development.**

**A)** Immunofluorescence staining of 100µm transverse vibratome sections of E14.5 jugular lymphatic structures from *Pde2a*$^{flox/flox}$*; Tie2-Cre* embryos (KO) and *Pde2a*$^{flox/flox}$ littermate embryos (CTRL) using antibodies against PROX1 (green) and Endomucin (red). Quantification of the jugular lymph sac (JLS) lumen from n = 4 KO and n = 4 CTRL embryos. Mean ± s.e.m., p-value: Unpaired Student's *t*-test. Scale bar: 500 µm. **B)** Immunofluorescence staining of 100µm transverse vibratome section of E13.5 jugular lymphatic structures from heterozygous *Pde2a*$^{flox/+}$*; Tie2-Cre* embryos using antibodies against PROX1 (green) and Endomucin (red). Single channel image for PROX1 is shown. Scale bar: 500 µm. **C)** Representative whole-mount immunofluorescence staining of the upper thoracic back skin of E14.5 *Pde2a*$^{-/-}$ and CTRL littermate embryos using an antibody against NRP2. Scale bar: 500 µm.

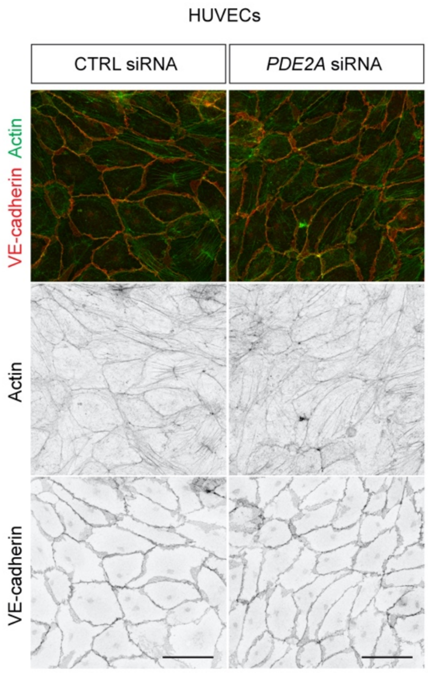


**Supplemental figure 4: PDE2A depletion selectively impairs lymphatic junctional stability.**
Representative immunofluorescence staining of 72h CTRL and *PDE2A* siRNA treated HUVECs using an antibody against VE-cadherin (red) and phalloidin (green, to visualize Actin). Single channel images are shown. Scale bar: 50 μm.

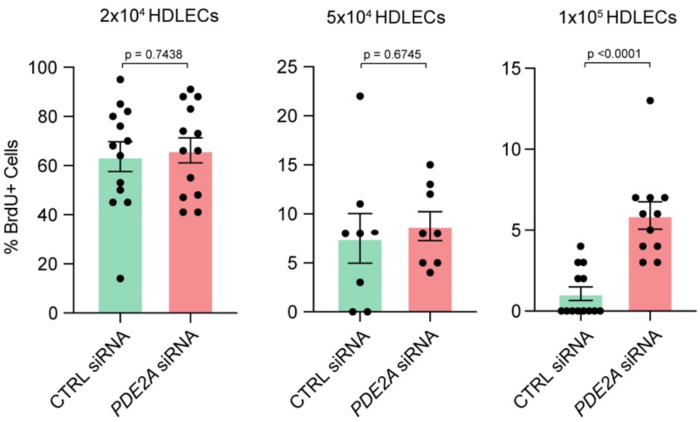


**Supplemental figure 5: Contact inhibition is prevented upon PDE2A depletion in high density lymphatic cell monolayers.**
Quantification of the percentage of BrdU$^+$ CTRL and *PDE2A* siRNA treated HDLECs (seeded at different cell densities) from n = 3 independent experiments at 2 x $10^4$ and 1 x $10^5$ cells per well, n = 2 independent experiments at 5 x $10^4$ cells per well. Mean ± s.e.m., p-value: Unpaired Student's *t*-test (with Welch's correction for 1 x $10^5$ cells per well).

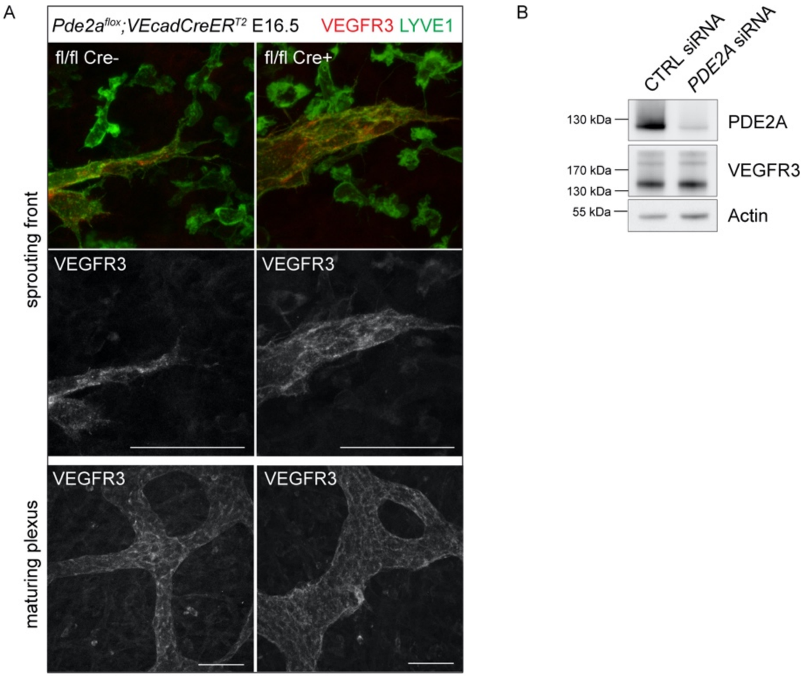


**Supplemental figure 6: VEGFR3 expression is not altered in the absence of PDE2A.**

**A)** Representative immunofluorescence staining of E16.5 whole-mount thoracic back skin sprouting front and maturing plexus from *Pde2a*$^{flox/flox}$*; Cdh5-CreER*$^{T2}$ embryos (KO) and *Pde2a*$^{flox/flox}$ embryos (CTRL) using antibodies against VEGFR3 (red) and LYVE1 (green). Single channel images for VEGFR3 are shown. Scale bar: 50 μm. **B)** Western Blot analysis of 72h CTRL and *PDE2A* siRNA-treated HDLECs. Lysates were probed with antibodies against PDE2A, VEGFR3 and Actin as loading control.

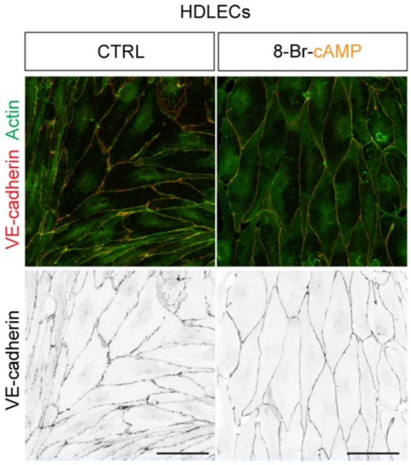


**Supplemental figure 7: cAMP stabilizes lymphatic endothelial junctions.**
Representative immunofluorescence staining of CTRL and 8-Br-cAMP treated HDLECs using an antibody against VE-cadherin (red) and phalloidin (green, to visualize Actin). Single channel images for VE-cadherin are shown. Scale bar: 50 μm.

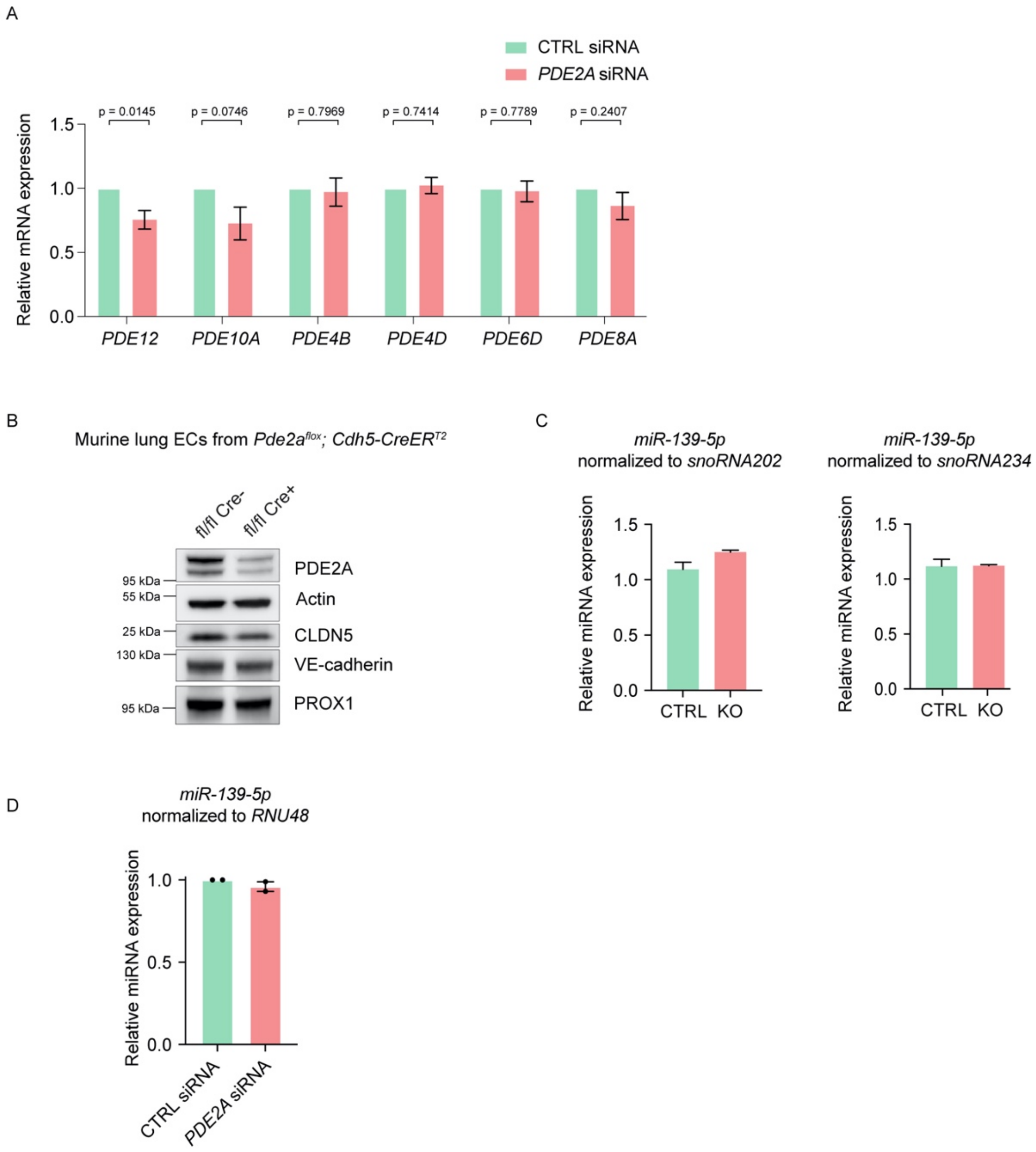


**Supplemental figure 8: Expression levels of lymphatic PDEs and *miR-139-5p* are not altered upon *PDE2A* siRNA treatment.**

**A)** Relative mRNA expression of lymphatic *PDE* transcripts in CTRL and *PDE2A* siRNA-treated HDLECs from n = 7 independent experiments. Mean ± s.e.m., p-value: One-sample *t*-test. **B)** Western Blot analysis of murine ECs isolated from 12-week-old *Pde2a*$^{flox/flox}$*; Cdh5-CreER*$^{T2}$ KO (n = 3) and *Pde2a*$^{flox/flox}$ littermate CTRL (n = 3) lungs. Cell lysates were probed with antibodies against PDE2A, CLDN5, VE-cadherin, PROX1 and Actin as loading control. **C)** Relative microRNA expression of *miR-139-5p* normalized to *snoRNA202* (left) and *snoRNA234* (right) in murine ECs isolated from KO (n = 3) and *Pde2a*$^{flox/flox}$ littermate CTRL (n = 3) lungs. **D)** Relative microRNA expression of *miR-139-5p* normalized to *RNU48* in CTRL and *PDE2A* siRNA-treated HDLECs from n = 2 independent experiments. Mean ± s.e.m., p-value
